# Cyclic Electromagnetic DNA Simulation (CEDS) can enhance the Functions of Anticancer miR-26a-5p and miR-126-5p in RAW 264.7 cells

**DOI:** 10.1101/2024.08.25.609548

**Authors:** Yeon Sook Kim, Suk Keun Lee

**Author notes:** Corresponding Author: Suk Keun Lee, DDS, MSD, PhD. Department of Oral Pathology, College of Dentistry, Gangneung-Wonju National University, 7, Jukheon-gil, Gangneung, 25457 South Korea.

## Abstract

Since cyclic electromagnetic DNA simulation (CEDS) was found to increase the hybridization potential of double-stranded DNA (dsDNA) and affect the functions of plasmid DNA (1), in this study, Dodecagonal CEDS was applied to RAW 264.7 cells to target anticancer miRNAs, miR-26a-5p and miR-126-5p, at 20-25 Gauss for 20 min during cell culture, followed by cytological observation, quantitative polymerase chain reaction (qPCR), and immunoprecipitation-based high-performance liquid chromatography (IP-HPLC). Both of miR-26a-5p*-CEDS and miR-126-5p*-CEDS induced significant decrease in cell number, and strong immunoreaction of c-caspase 3 in the remaining cells compared to oncogenic miR-21-5p*-CEDS and untreated control. In qPCR results, miR-26a-5p*-CEDS and miR-126-5p*-CEDS markedly increased the expression of target primary and mature miRNAs. IP-HPLC analysis showed that both miR-26a-5p*-CEDS and miR-126-5p*-CEDS induced potent anticancer effect on RAW 264.7 cells by suppressing RAS signaling, proliferation-growth-cytodifferentiation signaling axis, and angiogenesis-survival-oncogenesis-chronic inflammation axis compared to the untreated control and the positive control performed with non-specific poly-A 12A*-CEDS. MiR-26a-5p*-CEDS increased miRNA biogenesis and ER stress signaling more than miR-126-5p*-CEDS, while miR-126-5p*-CEDS increased NFkB signaling-immunity-apoptosis signaling axis more than miR-26a-5p*-CEDS. Therefore, it is proposed that the sequential treatment of miR-26a-5p*-CEDS and miR-126-5p*-CEDS may give a synergistic anticancer effect on RAW 264.7 cells. It is postulated that CEDS using a miRNA sequence can stimulate hybridization between miRNA and target mRNA and also increase the expression of target miRNA, and subsequently decrease the expression of miRNA target proteins, resulting in the alteration of protein signaling pathways in cells.

## Introduction

In the previous study, CEDS was able to target oligo-dsDNAs and plasmid DNAs, and resulted in the enhancement of hybridization and unique conformation of oligo-dsDNAs and the increase of RE digestion, *in vitro* RNA transcription, and production of green fluorescence protein (GFP), β-galactosidase, and ampicillin resistance protein from plasmid DNAs (*1*). Since dsDNAs may have different hybridization strengths and variable conformation depending on the buffer status as well as epigenetic modification, the targeting efficiency of CEDS may be uneventful in salt reaction buffer *in vitro* but may be possibly different in cellular cytoplasm or nuclear matrix. Therefore, in this study, dodecagonal CEDS targeting murine miRNAs in RAW 264.7 cells was performed.

CEDS was designed to provide electromagnetic stimulation to base pairs of dsDNA in a cyclic manner, targeting each base pair of dsDNA in decagonal or dodecagonal fashion at 20-25 Gauss. To achieve optimal electromagnetic induction, each A-T and G-C pair is stimulated with a different exposure time of 280 and 480 msec, respectively (*1*). This study employed both strands of target miRNAs in CEDS, which were designated as "target sequence*-CEDS."

### Cytological observation of RAW 264.7 cells treated with CEDS using a miRNA sequence

RAW 264.7 cells were cultured on cell culture slides (Hybridwell™, SPL, Korea) using the method described above. Approximately 50% confluent RAW 264.7 cells grown on culture slide surfaces were treated with dodecagonal CEDS using a murine miRNA sequence at 23-25 Gauss for 20 min and once more after 12 hours. After 24 hours culture, the cells were fixed with 10% buffered formalin, stained with hematoxylin and eosin, and underwent immunocytochemical (ICC) staining for c-caspase3.

ICC staining was performed using the indirect triple sandwich method on the Vectastatin system (Vector Laboratories, USA), and visualized using a 3-amino-9-ethylcarbazole solution (SantaCruz Biotechnology, USA) with no counter staining. The results were observed by optical microscope, and their characteristic images were captured and illustrated.

The negative control with no CEDS showed many cells diffusely scattered on the dish surface, which were occasionally positive for c-caspase3 (Fig 1 A-D), while the positive control treated with oncogenic miR-21-5p*-CEDS (TAGCTTATCAGACTGATGTTGA*-CEDS) showed overgrowth of cells that often aggregated into multicellular spheroids, which were weakly positive for c-caspase3 (Fig. 1 E-H). Whereas miR-26a-5p*-CEDS (TTCAAGTAATCCAGGATAGGCT*-CEDS) showed marked decrease in cell number, resulted in many empty spaces, and strong immunoreactivity for c-caspase3 in the remaining cells (Figure 1 I-L). The miR-126-5p*-CEDS (TTCAAGTAATCCAGGATAGGCT*-CEDS) also showed marked decrease in cell number, and the remaining cells were usually shrunken and strongly positive for c-caspase3 (Fig. 1 M-P). The cytological results demonstrated that miR-26a-5p* or miR-126-5p*-CEDS strongly inhibited proliferation and induced apoptosis of RAW 264.7 cells compared to untreated control and miR-21-5p*-CEDS.

**Figure 1.**
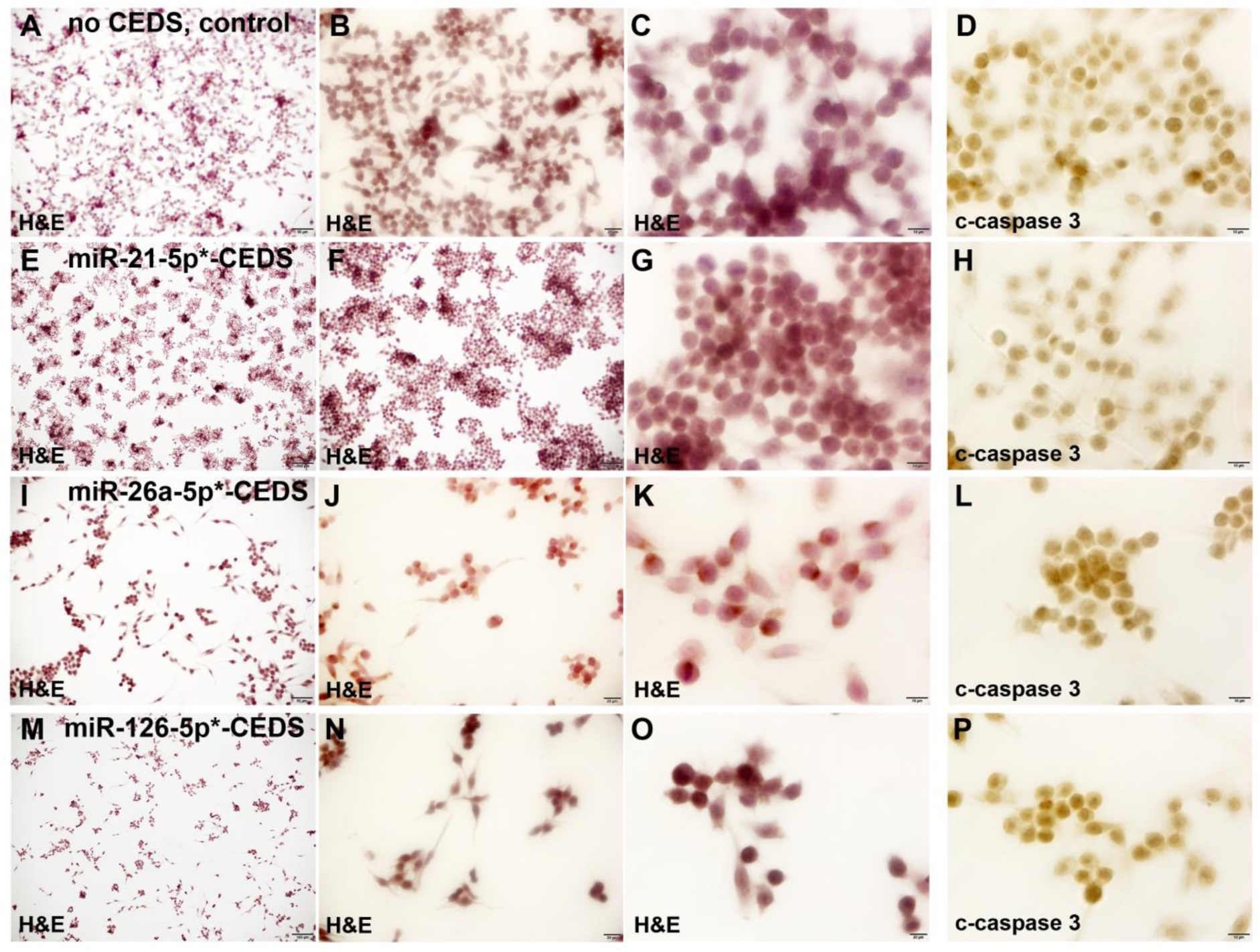
Cytological observation on RAW 264.7 cells treated with miR-26a-5p*-CEDS (I-L), miR-126-5p*-CEDS (M-P). miR-21-5p*-CEDS (positive control, E-H), or no CEDS (negative control, A-D). H&E staining and c-caspase3 immunostaining.

**Figure 2.**
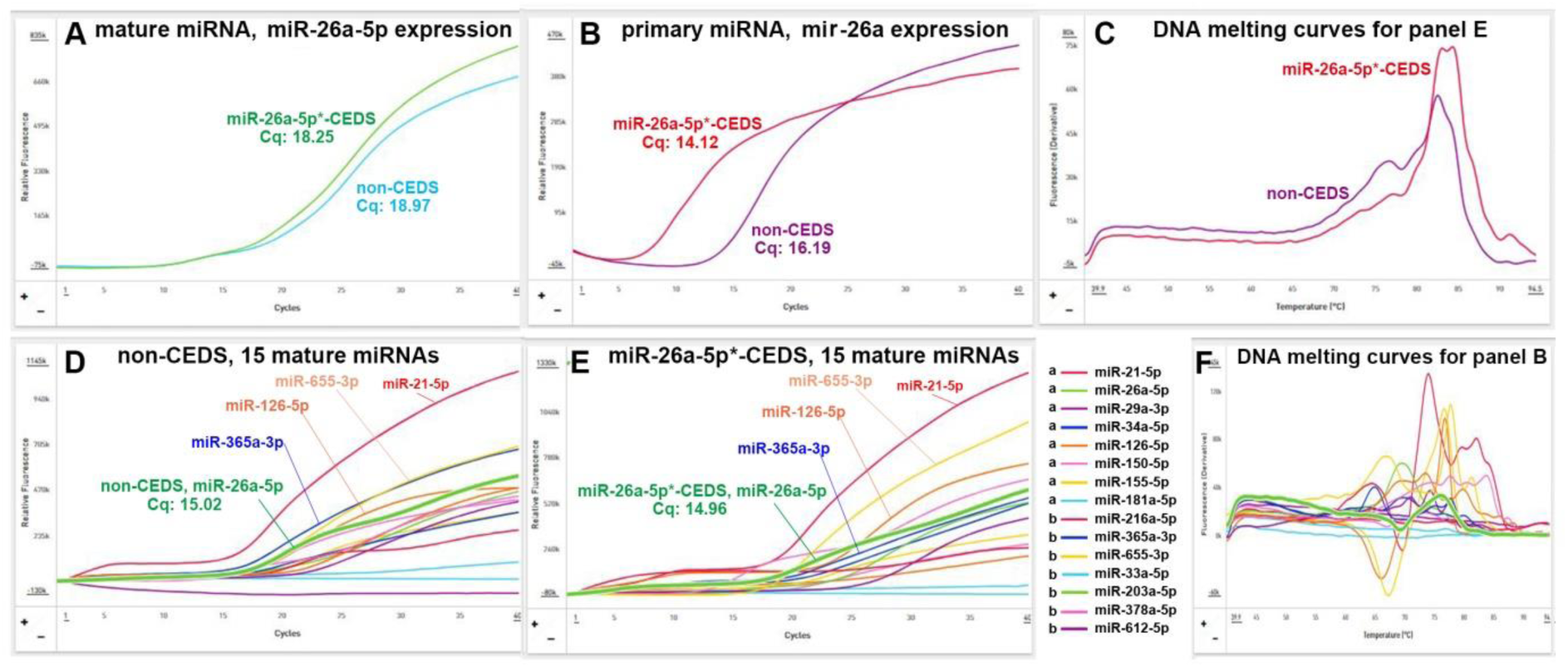
The effect of miR-26a-5p*-CEDS on the expression of mature miR-26a-5p and primary miR-26a in RAW 264.7 cells was evaluated by qPCR in comparison with other 14 miRNAs. *This figure represents three or more repeated experiments.

### MiR-26a-5p*-CEDS increased the expression of mature and primary miRNAs in qPCR analysis

For the quantitative polymerase chain reaction (qPCR), total miRNA was extracted from CEDS-treated cells using miRNeasy kit (Qiagen, USA), and reverse strand cDNA was generated using cDNA synthesis kit (CosmoGenetech, Korea). The cDNA was then used for qPCR (ChaiBio, USA) with a primer pair selected by DNA base pair polarity analysis of miRNA sequences (miRBase website) (Table S1). QPCR was carried out with varying annealing temperatures (43–53°C) and different PCR cycles (30–45 cycles) to obtain accurate Cq values, and followed by a DNA melting process to assess the fidelity of qPCR. QPCR was simultaneously conducted with other 14 miRNAs to compare their expression levels. The fold change of mature and primary miRNAs was calculated based on the Cq number.

The qPCR results showed that mmu-miR-26a-5p*-CEDS significantly increased the expression of mature miR-26a-5p and primary miR-26a, with average fold changes of 1.72 and 4.14, respectively, in comparison to untreated controls (Fig. 1 A,B). The multiple qPCRs for 15 miRNAs performed simultaneously showed that miR-26a-5p was moderately expressed compared to other miRNAs, and miR-26a-5p*-CEDS also affected the other miRNA expression by upregulating miR-126-5p and miR-655-3p while downregulating miR-365a-3p compared to untreated control (Figure 1 D,E), indicating the presence of secondary effect through the feedback loop interactions of gene transcription.

### IP-HPLC analysis for CEDS-induced changes of protein signaling pathways in RAW 264.7 cells

IP-HPLC has been developed to analyze the protein expression levels in comparison with the controls (*2, 3*). The immunoprecipitated proteins were subjected to the analysis with a HPLC (Agilent, USA) using a reverse phase column packed with non-adherent silica beads. The control and experimental samples were analyzed in sequence to permit a comparison of their HPLC peaks. IP-HPLC is available to use only small amount of total protein, up to 20-50 μg, thereby it can detect wide range of protein signaling simultaneously and repeatedly even with the limited amount of protein sample. However, IP-HPLC can provide the protein expression level in a simple, fast, accurate, cheap, multiple, and automatic way compared to other ordinary protein detection methods. In addition, the IP-HPLC data are available for statistical analysis.

The RAW 264.7 cells, derived from monocytes/macrophage-like cells of Balb/c mice, were utilized by limiting their culture passage number to less than 20 (*4*). The cells were cultured in Dulbecco’s modified Eagle’s medium supplemented with 10% fetal bovine serum, 100 units/mL carbenicillin, 100 μg/mL streptomycin, and 250 ng/mL amphotericin B (WelGene, Korea), in 5% CO_2_ incubator at 37.5°C. The cell culture product containing 10^8^-10^9^ cells was placed in the incubator and treated with dodecagonal CEDS using a target sequence at 20-25 Gauss, 5% CO2, and 37℃ for 20 minutes, left in the incubator for another 30 minutes, and harvested by centrifugation at 150g for 10 min for the following IP-HPLC assay.

The RAW 264.7 cells were lysed using protein lysis buffer (iNtRON Biotechnology, Korea), and analyzed by IP-HPLC as follow. Protein A/G agarose columns were separately pre-incubated with each 1 μg of 350 antibodies (Table S2). The supernatant of the antibody-incubated column was removed, and followed by IP-HPLC. Briefly, each protein sample, 50-100 μg, was mixed with 5 mL of binding buffer (150mM NaCl, 10mM Tris pH 7.4, 1mM EDTA, 1mM EGTA, 0.2mM sodium vanadate, 0.2mM PMSF and 0.5% NP-40) and incubated in the antibody-bound protein A/G agarose bead column on a rotating stirrer at room temperature for 1 h. After multiple washing of the columns with Tris-NaCl buffer, pH 7.5, in a graded NaCl concentration (0.15–0.3M), the target proteins were eluted with 300μL of IgG elution buffer (Pierce, USA). The immunoprecipitated proteins were analyzed using a precision HPLC unit (1100 series, Agilent, Santa Clara, CA, USA) equipped with a reverse-phase column and a micro-analytical UV detector system (SG Highteco, Hanam, Korea). Column elution was performed using 0.15M NaCl/20% acetonitrile solution at 0.5 mL/min for 15 min, 40°C, and the proteins were detected using a UV spectrometer at 280 nm. The control and experimental samples were run sequentially to allow comparisons.

For IP-HPLC, the whole protein peak areas (mAUs*) were obtained and calculated mathematically using an analytical algorithm by subtracting the negative control antibody peak areas, and protein expression levels were compared and normalized using the square roots of protein peak areas. IP-HPLC is available to use only small amount of total protein, up to 20-50 μg, thereby it can detect a wide range of protein signaling simultaneously and repeatedly even with the limited amount of protein sample. Since IP-HPLC gives the relative ratio (%) of protein expression compared to untreated control, the protein expression data could be analyzed statistically and widely compared with other protein expressions. The ratios were divided into five categories; severe underexpression (below 80%), marked underexpression (80%-below 95%), minimal change (95%-below 105%), marked overexpression (105%-120%), and severe overexpression (above 120%).

Although the immunoprecipitation is unable to define the size-dependent expression of target protein compared to western blot, it collects every protein containing a specific epitope against antibody in the protein samples from cells, blood, urine, saliva, inflammatory exudate, etc. (*2, 3, 5–14*). Therefore, IP-HPLC can detect whole precursor and modified target proteins similar to enzyme-linked immunosorbent assay (ELISA), but the antigen-antibody binding strength can be adjusted by different salt buffers depending on the condition of the protein samples.

### MiR-26a-5p*-CEDS induced potent anticancer protein signaling in RAW 264.7 cells

The RAW 264.7 cells treated with murine miR-26a-5p*-CEDS showed downregulation of 62 proteins (80.5%), upregulation of 11 proteins (14.3%), and minimal change of four proteins (5.2%) out of 77 miR-26a-5p target proteins, which were identified on the miRDB and TargetScan websites. In addition, miR-26a-5p*-CEDS indirectly downregulated 90 proteins (33%) and upregulated 93 proteins (34.1%) out of 273 miR-26a-5p non-target proteins simultaneously. The miR-26a-5p*-CEDS affected the overall protein signaling pathways in RAW 264.7 cells by downregulating miR-26a-5p target and reactive proteins as well (Fig. 3A, Table 1), which were summarized as follows according to the expression of miR-26a-5p target proteins.

**Figure 3.**
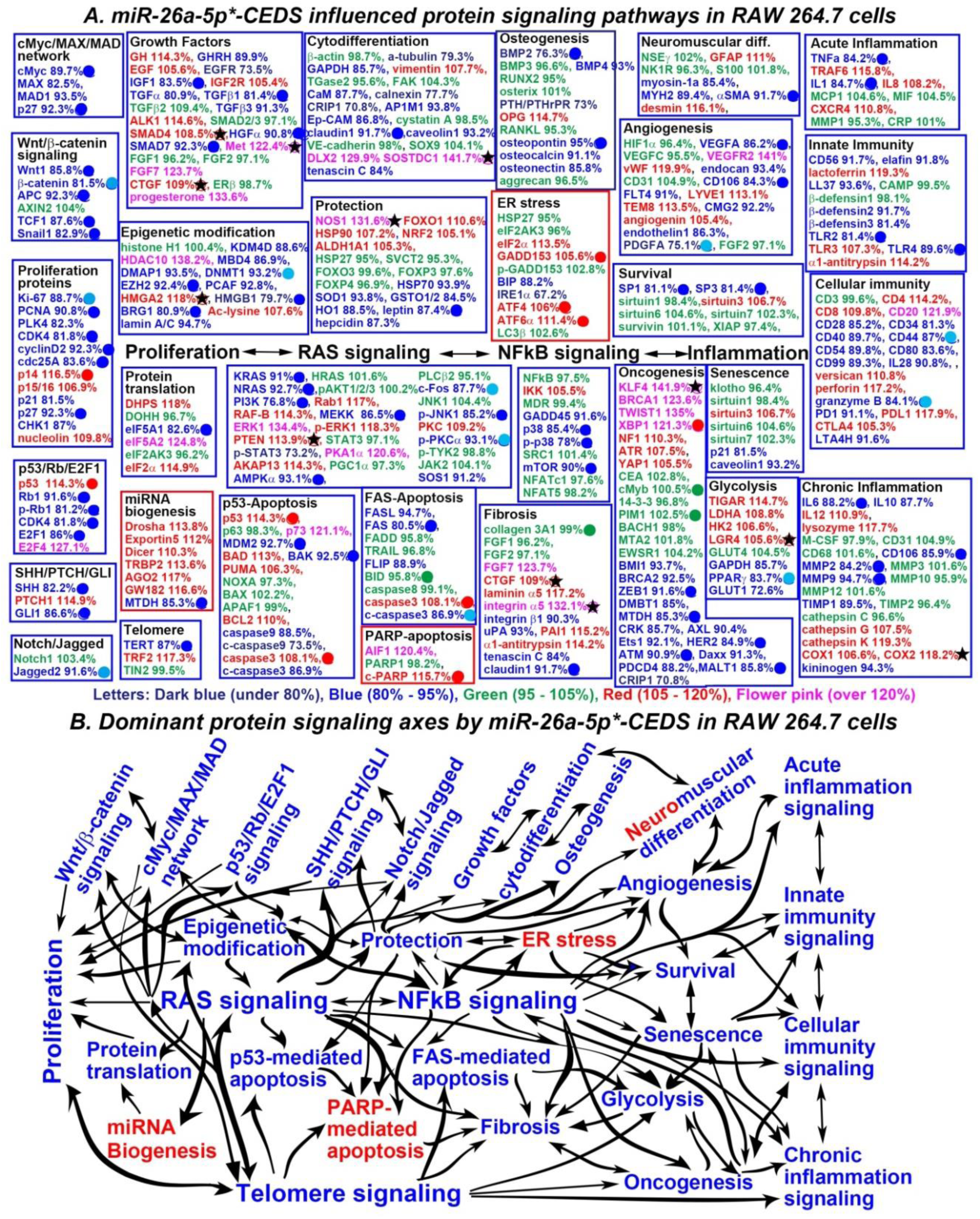
MiR-26a-5p*-CEDS influenced the protein signaling pathways (A) and axes (B) in RAW 264.7 cells. In IP-HPLC, proteins downregulated (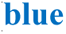), upregulated (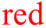), and minimally changed (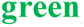) compared to untreated controls. Dominantly suppressed (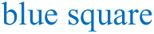) and activated (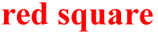) signaling. Downregulated proteins by direct targeting (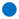) and indirect reactive (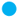), upregulated proteins by indirect response (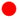), and minimal changed (±5%) proteins (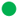). Some target proteins (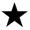) upregulated contrast to the current manuscripts and data from miRDB and TargetScan websites.

**Table 1.**
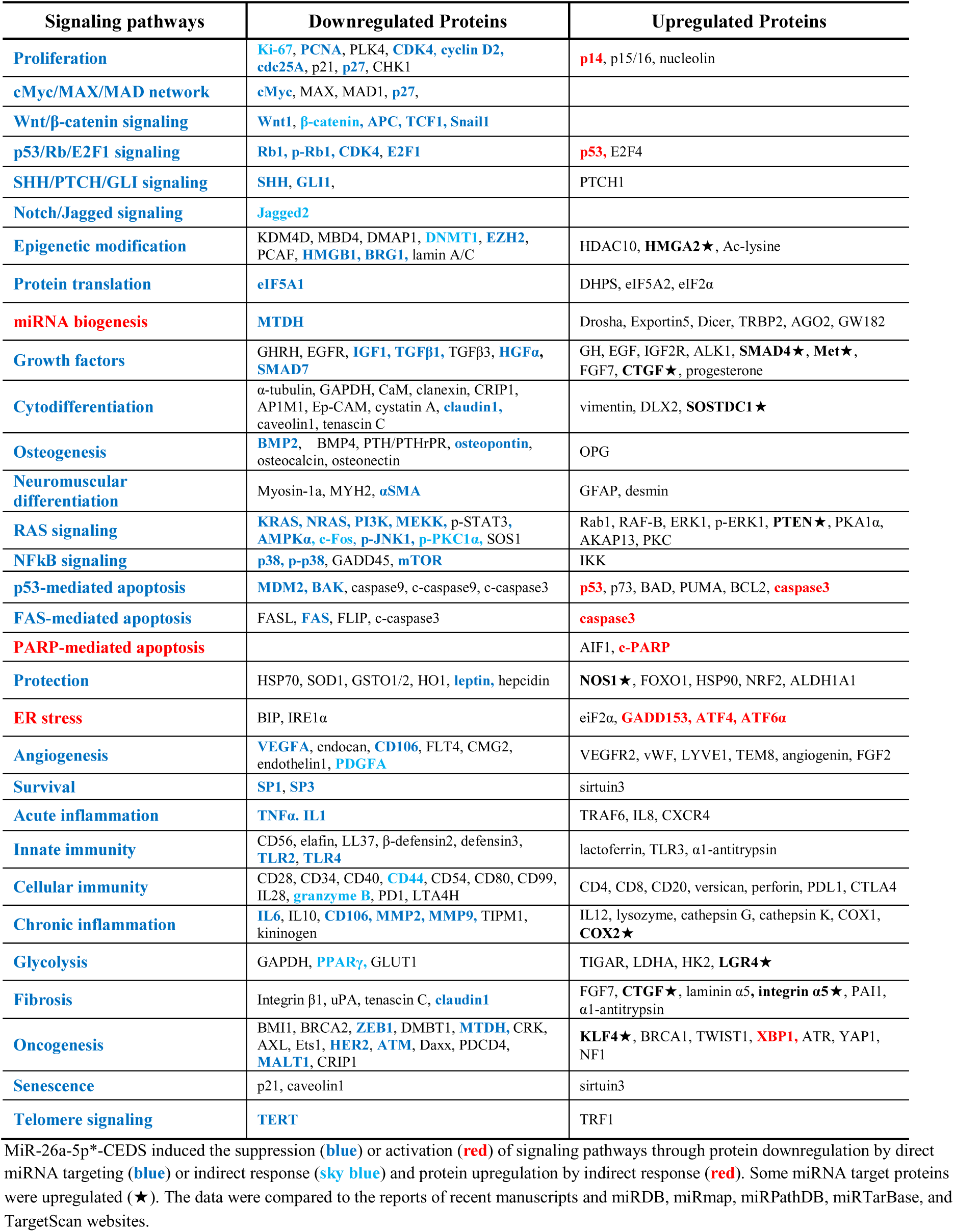
Protein expressions in RAW 264.7 cells treated with miR-26a-5p*-CEDS.

***Proliferation signaling*** was suppressed by downregulating Ki-67 (88.7%), PCNA (90.8%), CDK4 (81.8%), cyclin D2 (92.3%), cdc25A (83.6%), p27 (92. 3%). ***cMyc/MAX/MAD network*** was suppressed by downregulating cMyc (89.7%), p27 (92.3%). ***Wnt/β-catenin signaling*** was suppressed by downregulating Wnt1 (85.8%), β-catenin (81.5%), APC (92.3%), TCF1 (87.6%), Snail1 (82.9%).

***P53/Rb/E2F1 signaling*** was suppressed by downregulating Rb1 (91.6%), p-Rb (81.2%), CDK4 (81.8%), E2F1 (86%). ***SHH/PTCH/GLI signaling*** was suppressed by downregulating SHH (82.2%), GLI1 (86.6%).

***Notch/Jagged signaling*** was suppressed by downregulating Jagged2 (91.6%). ***Protein translation signaling*** appears to be suppressed by significantly downregulating eIF5A1 (82.6%). ***Growth factor signaling*** was suppressed by downregulating IGF1 (83.5%), TGFβ1 (81.4%), HGFα (90.8%), SMAD7 (92.3%).

***Cytodifferentiation signaling*** was suppressed by downregulating claudin1 (91.7%). ***Osteogenesis signaling*** was suppressed by downregulating BMP2 (76.3%), osteopontin (95%). ***Neuromuscular differentiation signaling*** appears to be suppressed by downregulating αSMA (91.7%). ***RAS signaling*** was suppressed by downregulating KRAS (91%), NRAS (92.7%), PI3K (76.8%), MEKK (86.5%), AMPKα (93.1%), c-Fos (87.7%), p-JNK1 (85.2%), SOS1 (91.2%).

***NFkB signaling*** was suppressed by downregulating p38 (85.4%), p-p38 (78%), mTOR (90%). ***Protection signaling*** was suppressed by downregulating leptin (87.4%). ***Angiogenesis signaling*** was suppressed by downregulating VEGFA (86.2%), CD106 (84.3%), PDGFA (75.1%). ***Survival signaling*** was suppressed by downregulating SP1 (81.1%), SP3 (81.4%).

***Acute inflammation signaling*** was suppressed by downregulating TNFα (84.2%), IL1 (84.7%). ***Innate immunity signaling*** was suppressed by downregulating TLR2 (81.4%), TLR4 (89.6%). ***Cellular immunity signaling*** was suppressed by downregulating CD44 (87%), granzyme B (84.1%). ***Chronic inflammation signaling*** was suppressed by downregulating IL6 (88.2%), CD106 (85.9%), MMP2 (84.2%), MMP9 (94.7%).

***P53-mediated apoptosis signaling*** was suppressed by downregulating MDM2 (92.7%), BAK (92.5%). ***FAS-mediated apoptosis signaling*** was inhibited by downregulating FAS (80.5%), c-caspase3 (86.9%). ***Fibrosis signaling*** was suppressed by downregulating claudin1 (91.7%). ***Senescence signaling*** was suppressed by downregulating p21 (81.5%), caveolin1 (93.2%). ***Glycolysis signaling*** was suppressed by downregulating PPARγ (83.7%), GLUT1 (72.6%). ***Telomere signaling*** was suppressed by downregulating TERT (87%).

Conversely, ***miRNA biogenesis signaling*** was activated by downregulating MTDH (85.3%). ***ER stress signaling*** was activated by upregulating GADD153 (105.6%), ATF4 (106%), ATF6α (111.4%). ***PARP-mediated apoptosis signaling*** was activated by upregulating c-PARP (115.7%).

In addition, miR-26a-5p*-CEDS markedly inactivated ***epigenetic modification signaling*** by downregulating DNMT1 (93.2%), EZH2 (92.4%), HMGB1 (79.7%), BRG1 (80.9%). MiR-26a-5p*-CEDS significantly suppressed ***oncogenesis signaling*** by downregulating ZEB1 (91.6%), MTDH (85.3%), HER2 (84.9%), ATM (90.9%), MALT1 (85.8%). Besides the expressions of miR-26a-5p target and reactive proteins, the expressions of miR-26a-5p non-target proteins were summarized in Table 1.

Consequently, miR-26a-5p*-CEDS significantly impacted the entire protein signaling pathways in RAW 264.7 cells. MiR-26a-5p*-CEDS markedly suppressed ***RAS and NFkB signaling axis***, leading to the suppression of ***proliferation-related signaling axis, protection-survival-senescence signaling axis, inflammation-angiogenesis-fibrosis signaling axis, oncogenesis-glycolysis signaling axis, telomere-epigenetic modification signaling axis,*** and ***p53-and FAS-mediated apoptosis signaling axis***. Conversely, it activated ***miRNA biogenesis signaling, ER stress signaling,*** and ***PARP-mediated apoptosis signaling*** (Fig. 3B).

The results indicate that miR-26a-5p*-CEDS may provide a potent anticancer effect on RAW 264.7 cells and consistently exert apoptotic cell death by targeting 62 proteins (80.5%) out of 77 miR-26a-5p target proteins. Conversely, it may attenuate the innate and cellular immunity and cytodifferentiation, which are essential for immune surveillance and cellular maturation.

### MiR-126-5p*-CEDS increased the expression of mature and primary miRNAs in qPCR analysis

The qPCR results showed that mmu-miR-126-5p*-CEDS increased the expression of mature miR-126-5p and primary miR-126, with average fold changes of 3.36 and 2.56, respectively, in comparison to untreated controls (Fig. 4 A,B). The multiple qPCRs for 15 miRNAs performed simultaneously showed that miR-126-5p was moderately expressed compared to other miRNAs, and miR-126-5p*-CEDS also affected the other miRNA expression by upregulating miR-26a-5p and miR-655-3p compared to untreated control (Fig. 4 D,E), indicating the presence of secondary effect through the feedback loop interactions of gene transcription.

**Figure 4.**
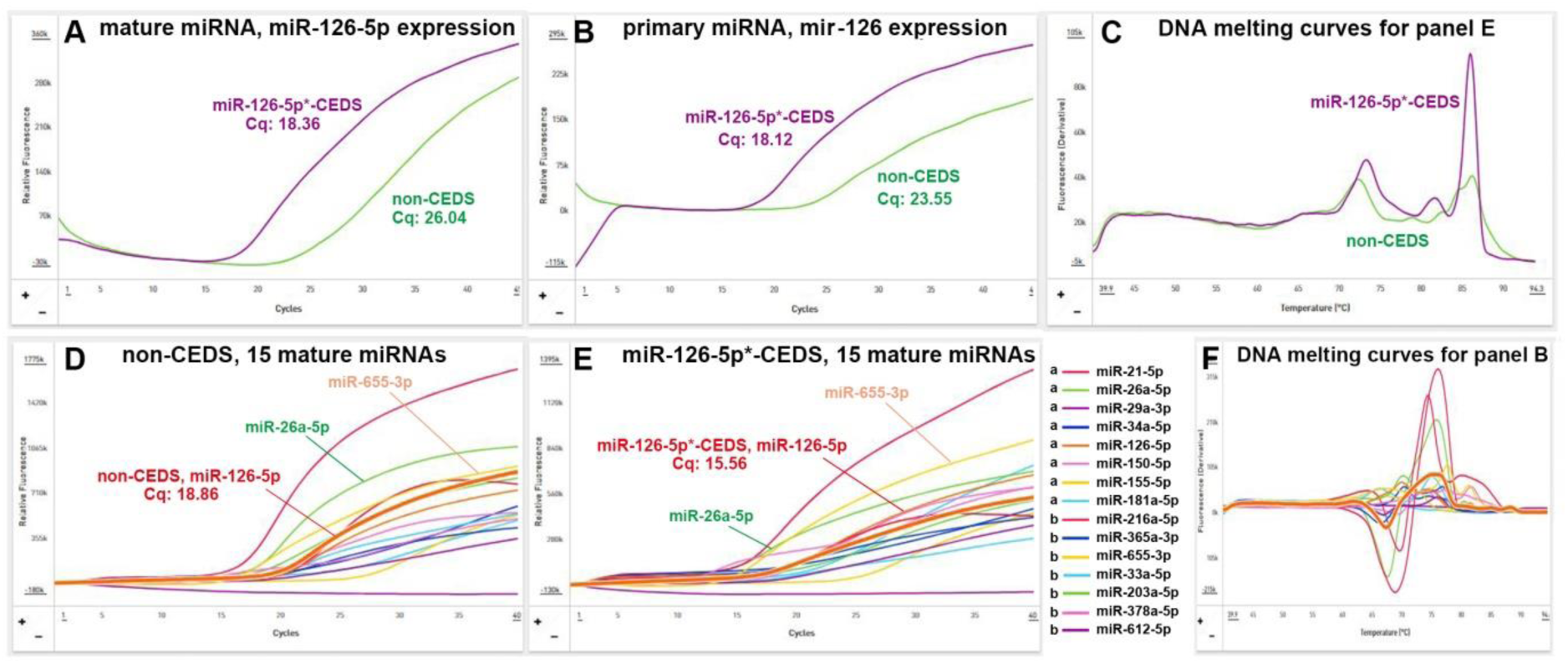
The effect of miR-126-5p*-CEDS on the expression of mature miR-126-5p and primary miR-126 in RAW 264.7 cells was evaluated by qPCR in comparison with other 14 miRNAs. *This figure represents three or more repeated experiments.

### MiR-126-5p*-CEDS induced significant anticancer signaling in RAW 264.7 cells

The RAW 264.7 cells treated with miR-126-5p*-CEDS showed downregulation of 46 proteins (57.5%), upregulation of 13 proteins (16.3%), and minimal change of 20 proteins (25%) out of 80 miR-126-5p target proteins, which were identified on the miRDB and TargetScan websites. In addition, miR-126-5p*-CEDS indirectly downregulated 98 proteins (36.3%) and upregulated 95 proteins (35.2%) out of 270 miR-126-5p non-target proteins simultaneously. The miR-126-5p*-CEDS affected the overall protein signaling pathways in RAW 264.7 cells by downregulating miR-126-5p target and reactive proteins as well (Fig. 5A, Table 2), which were summarized as follows according to the expression of miR-126-5p target proteins.

**Figure 5.**
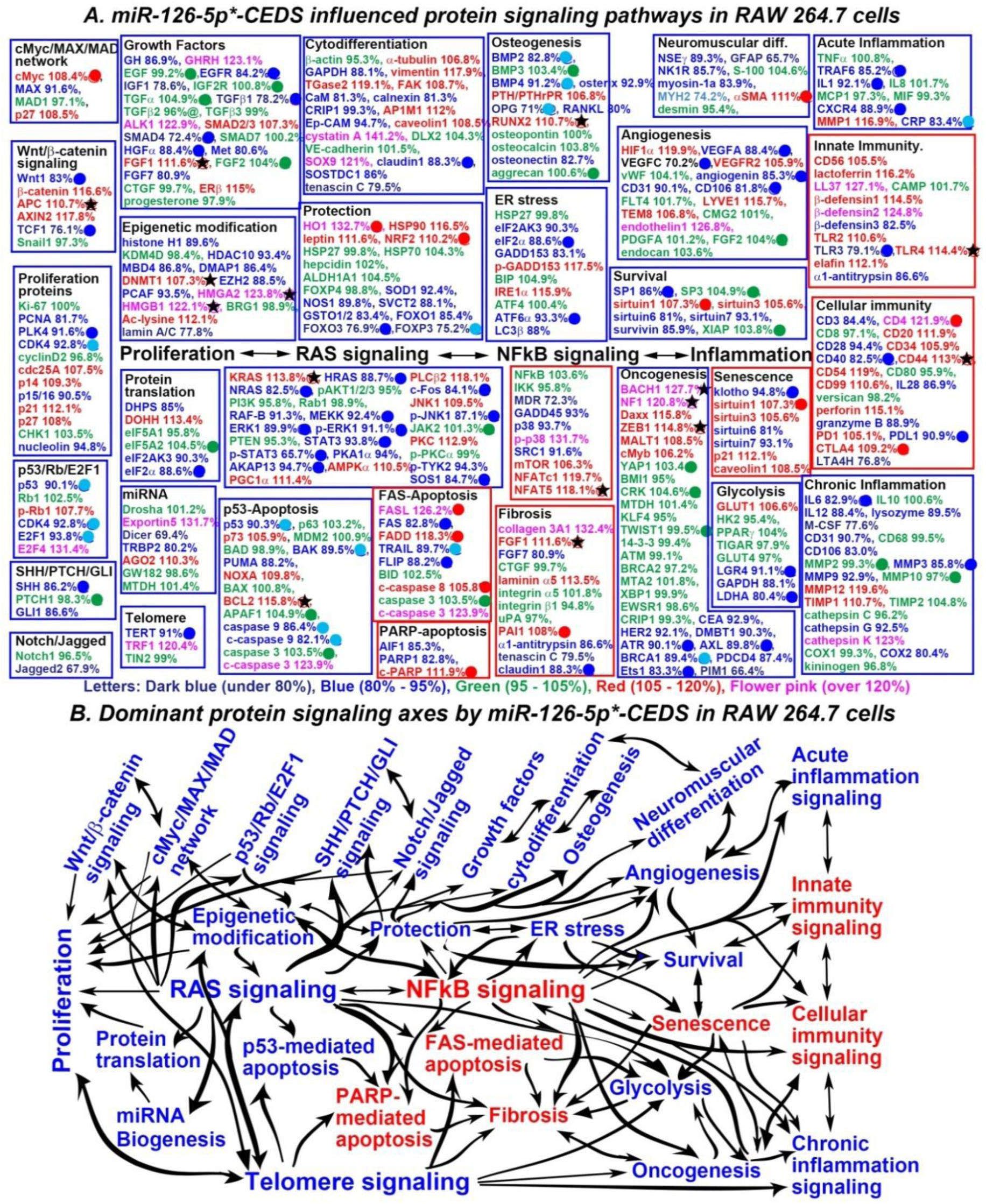
MiR-126-5p*-CEDS influenced the protein signaling pathways (A) and axes (B) in RAW 264.7 cells. In IP-HPLC, proteins downregulated (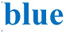), upregulated (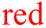), and minimally changed (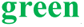) compared to untreated controls. Dominantly suppressed (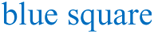) and activated (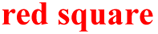) signaling. Downregulated proteins by direct targeting (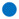) and indirect reactive (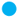), upregulated proteins by indirect response (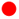), and minimal changed (±5%) proteins (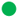). Some target proteins (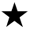) upregulated contrast to the current manuscripts and data from miRDB and TargetScan websites.

**Table 2.** Protein expressions in RAW 264.7 cells treated with miR-126-5p*-CEDS.

| Signaling pathways | Downregulated Proteins | Upregulated Proteins |
| --- | --- | --- |
| Proliferation | PCNA, **PLK4**, **CDK4**, p15/16, nucleolin | cdc25A, p14, p21, p27 |
| cMyc/MAX/MAD network | MAX | **cMyc**, p27 |
| Wnt/β-catenin signaling | **Wnt1**, **TCF1** | β-catenin, APC, AXIN2 |
| p53/Rb/E2F1 signaling | **p53**, **CDK4**, **E2F1** | p-Rb1, E2F4 |
| SHH/PTCH/GLI signaling | **SHH**, GLI1 |  |
| Notch/Jagged signaling | Jagged |  |
| Epigenetic modification | Histone H, HDAC10, MBD4, DMAP, EZH2, PCAF, lamin A/C | **DNMT1**★, **HMGA2**★, **HMGB1**★, Ac-lysine |
| Protein translation | DHPS, eIF2AK3, **eIF2α** |  |
| miRNA biogenesis | Dicer, TRBP2 | Exportin5, AGO2 |
| Growth factors | GH, **EGFR**, IGF1, **TGFβ1**, **SMAD4**, **HGFα**, Met, FGF7 | GHRH, TGFα, ALK1, SMAD2/3, **FGF1**★, FGF2, ERβ |
| Cytodifferentiation | GAPDH, CaM, calnexin, CRIP1, Ep-CAM, **claudin1**, SOSTDC1, tenascin C | α-tubulin, vimentin, TGase2, FAK, AP1M1, caveolin1, cystatin A, SOX9 |
| Osteogenesis | **BMP2**, **BMP4**, osterix, **OPG**, RANKL, osteonectin | **RUNX2**★, PTH/PTHrPR |
| Neuromuscular differentiation | NSEγ, GFAP, NK1R, myosin-1a | **αSMA** |
| RAS signaling | **HRAS**, **NRAS**, RAF-B, **MEKK**, **ERK1**, **p-ERK1**, **STAT3**, **p-STAT3**, PKA1α, AKAP13, **c-Fos**, **p-JNK1**, p-TYK2, **SOS1** | **KRAS**★, AMPKα, PGC1α, PLCβ2, JNK1, PKC |
| NFκB signaling | MDR, GADD45, p38, SRC1 | p-p38, mTOR, NFATc1, **NFAT5**★ |
| p53-mediated apoptosis | **p53**, **BAK**, **PUMA**, **caspase9**, **c-caspase9** | p73, NOXA, **BCL2**★, APAF1, c-caspase3 |
| FAS-mediated apoptosis | **FAS**, **TRAIL**, **FLIP** | **FASL**, **FADD**, **c-caspase8**, c-caspase3 |
| PARP-mediated apoptosis | AIF1, PARP1 | **c-PARP** |
| Protection | SOD1, NOS1, SVCT2, GSTO1/2, FOXO1, **FOXO3**, **FOXP3** | **HO1**, HSP90, leptin, **NRF2** |
| ER stress | eIF2AK3, **eIF2α**, GADD153, **ATF6α**, LC3β | p-GADD153, IRE1α |
| Angiogenesis | **VEGFA**, **VEGFC**, **angiogenin**, CD31, **CD106** | HIF1α, VEGFR2, LYVE1, TEM8, endothelin1, FGF2 |
| Survival | **SP1**, sirtuin6, sirtuin7, survivin, | SP3, **sirtuin1**, sirtuin3, XIAP |
| Acute inflammation | **TRAF6**, **IL1**, **CXCR4**, **CRP** | MMP1 |
| Innate immunity | β-defensin3, **TLR3**, α1-antitrypsin | CD56, lactoferrin, LL37, β-defensin1, β-defensin2, TLR2, **TLR4**★, elafin |
| Cellular immunity | CD3, CD28, **CD40**, IL28, granzyme B, **PDL1**, LTA4H | **CD4**, CD20, CD34, **CD44**★, CD54, CD99, perforin, PD1, **CTLA4** |
| Chronic inflammation | **IL6**, IL12, lysozyme, M-CSF, CD31, CD106, **MMP3**, MMP9, cathepsin G, COX2 | MMP12, TIMP1, cathepsin K |
| Glycolysis | **LGR4**, GAPDH, **LDHA** | GLUT1 |
| Fibrosis | FGF7, α1-antitrypsin, tenascin, **claudin1** | collagen 3A1, FGF1, laminin α5, **PAI1** |
| Oncogenesis | CEA, HER2, DMBT1, **ATR**, **AXL**, **BRCA1**, PDCD4, **Ets1**, PIM1 | **BACH1**★, **NF1**★, Daxx, **ZEB1**★, MALT1, YAP1, cMyb |
| Senescence | **klotho**, sirtuin6, sirtuin7 | **sirtuin1**, sirtuin3, p21, caveolin1 |
| Telomere signaling | **TERT** | TRF1 |

MiR-126-5p\*-CEDS induced the suppression (blue) or activation (red) of signaling pathways through protein downregulation by direct miRNA targeting (blue) or indirect response (sky blue) and protein upregulation by indirect response (red). Some miRNA target proteins were upregulated (★). The data were compared to the reports of recent manuscripts and miRDB, miRmap, miRPathDB,

***Proliferation signaling*** was suppressed by downregulating PLK4 (91.6%), CDK4 (92.8%). cMyc/MAX/MAD network was suppressed by downregulating MAX (91.6%). ***Wnt/β-catenin signaling*** was suppressed by downregulating Wnt1 (83%), TCF1 (76.1%).

***P53/Rb/E2F1 signaling*** was suppressed by downregulating p53 (90.1%), CDK4 (92.8%), E2F1 (93.8%). ***SHH/PTCH/GLI signaling*** was suppressed by downregulating SHH (86.2%). ***Notch/Jagged signaling*** was suppressed by downregulating Jagged2 (67.9%). ***Protein translation signaling*** was suppressed by downregulating DHPS (85%), eIF2α (88.6%).

***MiRNA biogenesis signaling*** appears to be inactivated by downregulating Dicer (69.4%), TRBP2 (80.2%). ***Growth factor signaling*** was suppressed by downregulating EGFR (84.2%), TGFβ1 (78.2%), SMAD4 (72.4%), HGFα (88.4%). ***Cytodifferentiation signaling*** appears to be inactivated by downregulating claudin1 (88.3%). ***Osteogenesis signaling*** was suppressed by downregulating BMP2 (82.8%), BMP4 (91.2%), OPG (71%).

***Neuromuscular differentiation signaling*** was inactivated by downregulating NSEγ (89.3%), GFAP (65.7%), NK1R (85.7%), myosin-1a (83.9%). ***RAS signaling*** was suppressed by downregulating HRAS (88.7%), NRAS (82.5%), MEKK (92.4%), ERK1 (89.9%), p-ERK1 (91.1%), STAT3 (93.8%), p-STAT3 (65.7%), AKAP13 (94.7%), c-Fos (84.1%), p-JNK1 (87.1%), SOS1 (84.7%). ***Angiogenesis signaling*** was suppressed by downregulating VEGFA (88.4%), VEGFC (70.2%), angiogenin (85.3%), CD106 (81.8%). ***Survival signaling*** was suppressed by downregulating SP1 (86%), survivin (85.9%).

***Acute inflammation signaling*** was suppressed by downregulating TRAF6 (85.2%), IL1 (84.7%), CXCR4 (88.9%). ***Chronic inflammation signaling*** was suppressed by downregulating IL6 (82.9%), MMP3 (85.8%), COX2 (80.4%). ***p53-mediated apoptosis signaling*** was suppressed by downregulating p53 (90.3%), BAK (89.5%), caspase9 (86.4%), c-caspase9 (82.1%). ***Glycolysis signaling*** was suppressed by downregulating LGR4 (91.1%), LDHA (80.4%). ***Telomere signaling*** was suppressed by downregulating TERT (91%).

Conversely, ***innate immunity signaling*** was activated by upregulating CD56 (105.5%), lactoferrin (116.2%), LL37 (127.1%), β-defensin1 (114.5%), β-defensin2 (124.8%), TLR2 (110.6%), TLR4 (114.4%). ***Cellular immunity signaling*** was activated by upregulating CD4 (121.9%), CTLA4 (109.2%). ***NFkB signaling*** was activated by upregulating p-p38 (131.7%), mTOR (106.3%), NFATc1 (119.7%), NFAT5 (118.1%).

***FAS-mediated apoptosis signaling*** was activated by upregulating FASL (126.2%), FADD (118.3%), c-caspase8 (105.8%). ***PARP-mediated apoptosis signaling*** was activated by upregulating c-PARP (111.9%). ***Fibrosis signaling*** was activated by upregulating PAI1 (108%). ***Senescence signaling*** appears to be activated by upregulating sirtuin1 (107.3%).

In addition, miR-126-5p*-CEDS variably influenced ***epigenetic modification signaling*** by downregulating histone H1 (89.6%), HDAC10 (93.4%), MBD4 (86.8%), DMAP1 (86.4%), EZH2 (88.5%), PCAF (93.5%), lamin A/C (77.8%). MiR-126-5p*-CEDS broadly affected ***oncogenesis signaling*** by downregulating ATR (90.1%), AXL (89.8%), BRCA1 (89.4%), Ets1 (83.3%). Besides the expressions of miR-126-5p target and reactive proteins, the expressions of miR-126-5p non-target proteins were summarized in Table 2.

Consequently, miR-126-5p*-CEDS significantly impacted the entire protein signaling pathways in RAW 264.7 cells. MiR-126-5p*-CEDS suppressed ***RAS signaling pathway***, resulting in the inactivation of ***proliferation-related signaling axis***, ***protection-ER stress-angiogenesis-survival signaling axis, glycolysis-oncogenesis-chronic inflammation signaling axis,*** and ***telomere-epigenetic modification-RAS-p53 mediated apoptosis signaling axis***. Conversely, it enhanced ***NF-kB signaling***, which led to the activation of ***FAS-and PARP-mediated apoptosis-fibrosis signaling axis*** and ***senescence-innate and cellular immunity signaling axis*** (Fig. 5B).

The results indicate that miR-126-5p*-CEDS induced a strong anticancer effect on RAW 264.7 cells by suppressing proliferation, angiogenesis, survival, glycolysis, chronic inflammation, and telomere instability, concomitantly enhancing FAS-and PARP-mediated apoptosis and innate and cellular immunity with targeting 46 proteins (57.5%) out of 80 miR-126-5p target proteins. It is also noteworthy that miR-126-5p*-CEDS may induce senescence and fibrosis of cells as a result.

### Nonspecific poly-A 12A*-CEDS induced imbalanced protein signaling in RAW 264.7 cells

RAW 264.7 cells were also treated with CEDS using nonspecific poly-A sequence (12A*-CEDS) as a positive control, and analyzed by IP-HPLC to compared with the results of miR-26a-5p*-CEDS and miR-126-5p*-CEDS. In contrast, 12A*-CEDS resulted in imbalanced protein signaling in RAW 264.7 cells, activating epigenetic methylation, oncogenesis, and telomere instability, leading to chronic inflammation and apoptosis, in the absence of cellular growth and differentiation, wound healing, and ROS protection (Fig. 6, Table 4). The variable protein expressions after 12A*-CEDS were detected by IP-HPLC using 350 antisera, and their imbalaced influences in different signaling pathways were summarized and described in Supplement Text. Therefore, it is postulated that 12A*-CEDS induced irregular upregulation and downregulation of multiple signaling pathways due to the irregular stimulation of gene transcription by targeting poly-A sequences widely distributed in genome, resulted in the aging and retrogressive changes with a high potential of oncogenesis.

**Figure 6.**
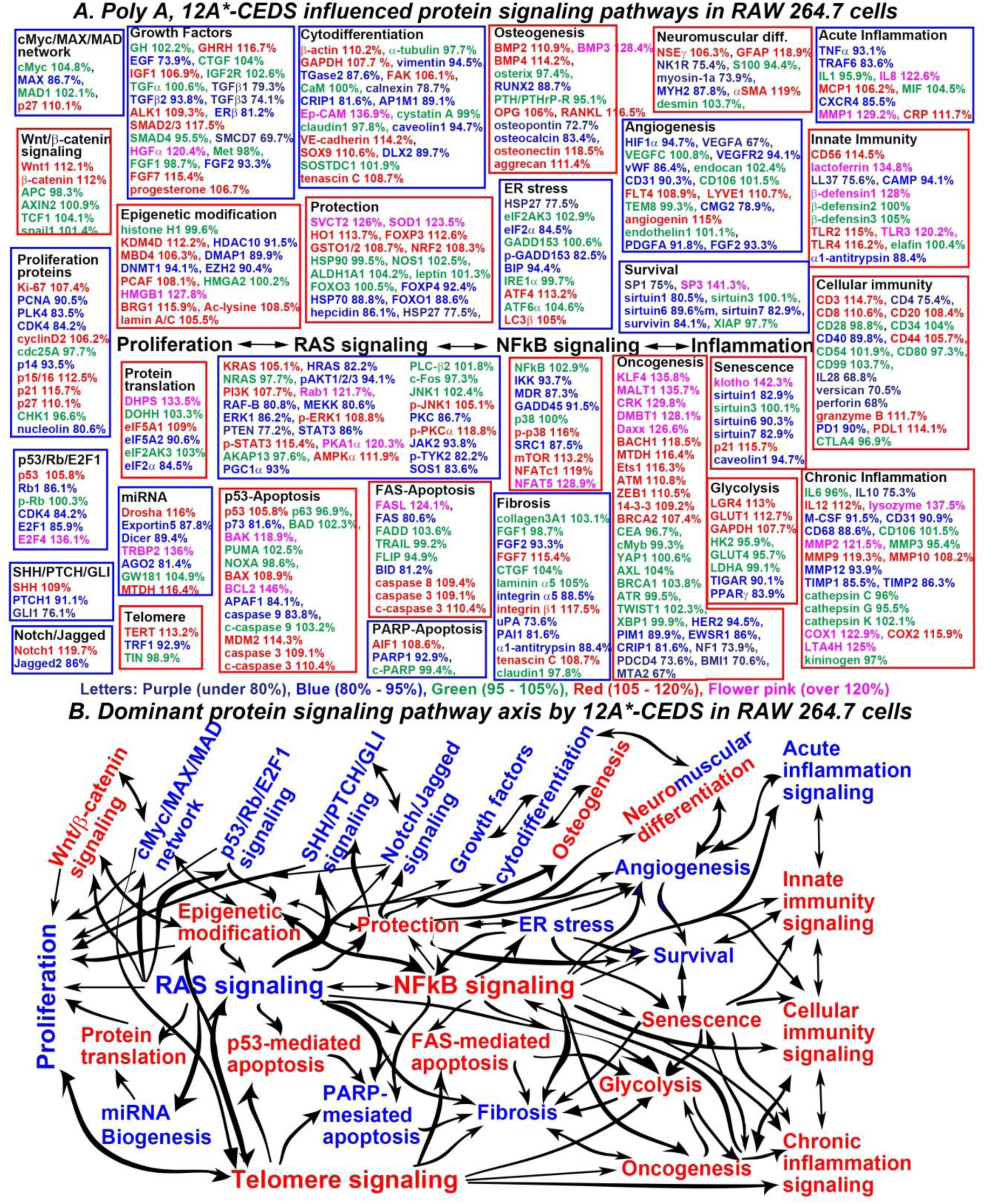
The nonspecific sequence 12A*-CEDS influenced the protein signaling pathways (A) and axis (B) in RAW 264.7 cells. The IP-HPLC results revealed proteins that were downregulated (blue), upregulated (red), and minimally changed (green) compared to the untreated controls. This allowed the dominant trend of suppressed (blue square) and activated (red square) signaling pathways to be defined.

## Discussion

The human genome encodes approximately 2,600 mature miRNAs (miRBase V.22), and it is known that a single miRNA can modulate thousands of genes by recognizing complementary sequences not only in the 3’ UTR region, but also in the 5’ UTR and open reading frame regions (*15*). In this study, miR-26a-5p*-CEDS downregulated 80.5% of target proteins (62/77) and simultaneously 33% of non-target proteins (90/273%), and miR-126-5p*-CEDS downregulated 57.5% of target proteins (46/80) and simultaneously 36.3% of non-target proteins (98/270) (Table 3). Moreover, the CEDS increased the expression of targeting miRNA together with other miRNAs, such as miR-655-3p, in a synergistic manner.

**Table 3.**
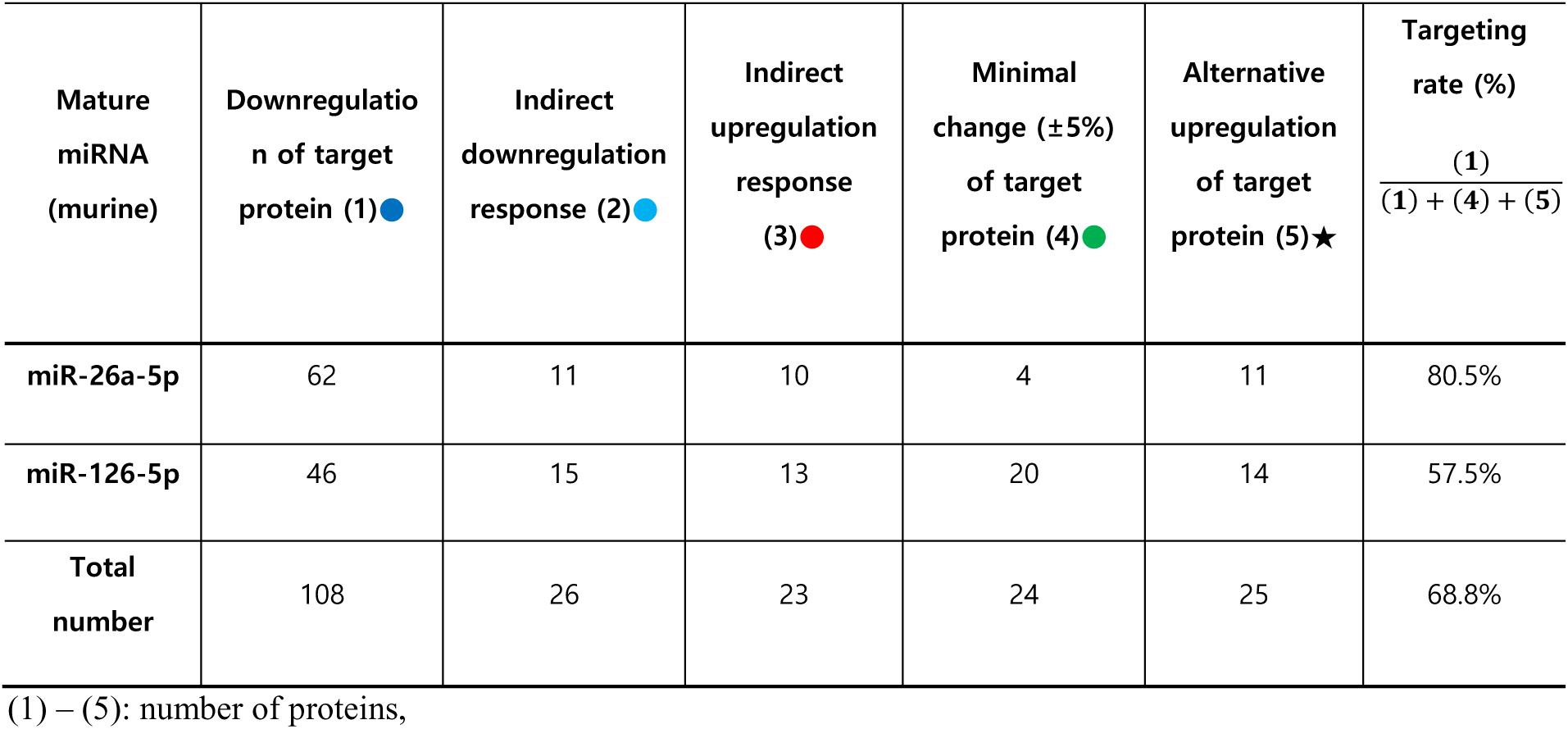
The efficiency of CEDS-induced miRNA targeting in the protein expression (n=350) of RAW 264.7 cells.

**Table 4.** Protein expressions in RAW 264.7 cells treated with nonspecific sequence (12A)*-CEDS.

| Signaling pathways | Downregulated Proteins | Upregulated Proteins |
| --- | --- | --- |
| Proliferation | PCNA, PLK4, CDK4, p14, nucleolin | Ki-67, cyclin D2, p15/16, p21, p27 |
| cMyc/MAX/MAD network | MAX | p27 |
| Wnt/β-catenin signaling | | Wnt1, β-catenin |
| p53/Rb/E2F1 signaling | Rb1, CDK4, E2F1 | p53, E2F4 |
| SHH/PTCH/GLI signaling | PTCH1, GLI1 | SHH |
| Notch/Jagged signaling | Jagged2 | Notch1 |
| Epigenetic modification | HDAC10, DMAP1, DNMT1, EZH2 | KDM4D, MBD4, PCAF, HMGB1, BRG1, Ac-lysine, lamin A/C |
| Protein translation | eIF5A2, eIF2α | DHPS, eIF5A1 |
| miRNA biogenesis | Exportin5, Dicer, AGO2, | Drosha, TRBP, MTDH |
| Growth factors | EGF, TGFβ1, TGFβ2, TGFβ3, SMCD7, FGF2, ERβ | GHRH, NF1, ALK1, SMCD2/3, HGFα, FGF7, progesterone |
| Cytodifferentiation | Vimentin, TGase2, calnexin, CRIP1, AP1M1, caveolin1, DLX2 | β-actin, GAPDH, FAK, Ep-CAM, VE-cadherin, SOX9, tenascin C |
| Osteogenesis | RUNX2, osteopontin, osteocalcin, | BMP2, BMP3, BMP4, OPG, RANKL, osteonectin, aggrecan |
| Neuromuscular differentiation | NK1R, myosin-1a, MYH2 | NSEγ, GFAP, αSMA |
| RAS signaling | HRAS, pAKT1/2/3, RAF-B, MEKK, ERK1, PTEN, STAT3, PGC1α, PKC, JAK2, p-TYK2, SOS1 | KRAS, PI3K, Rab1, p-STAT3, PKA1α, AMPKα, p-JNK1, p-PKC1α |
| NFκB signaling | IKK, MDR, GADD45, SRC1 | p-p38, mTOR, NFATc1, NFAT5 |
| p53-mediated apoptosis | p73, APAF1, caspase9 | p53, BAK, BAX, BCL2, MDM2, caspase3, c-caspase3 |
| FAS-mediated apoptosis | FAS, BID | FASL, caspase8, caspase3, c-caspase3 |
| PARP-mediated apoptosis | PARP1 | AIF1 |
| Protection | FOXP4, HSP70, FOXO1, hepcidin, HSP27 | SVCT2, SOD1, HO1, FOXP3, GSTO1/2, NRF2, |
| ER stress | HSP27, eIF2α, p-GADD153, BIP | ATF4, LC3β |
| Angiogenesis | HIF1α, VEGFA, VEGFR2, vWF, CD31, CMG2, PDGFA, FGF2 | FLT4, LYVE1, angiogenin |
| Survival | SP1, sirtuin1, sirtuin1, sirtuin6, sirtuin7, survivin | SP3 |
| Acute inflammation | TNFα, TRAF6, CXCR4 | IL8, MCP1, MMP1, CRP |
| Innate immunity | LL37, CAMP, α1-antitrypsin | CD56, lactoferrin, TLR2, TLR3, TLR4 |
| Cellular immunity | CD4, CD40, IL28, versican, perforin, PD1 | CD3, CD8, CD20, CD44, granzyme B, PDL1 |
| Chronic inflammation | IL6, IL10, M-CSF, CD31, CD68, MMP12, TIMP1, TIMP2 | IL12, lysozyme, MMP2, MMP9, MMP10, COX1, COX2, LTA4H |
| Glycolysis | TIGAR, PPARγ | LGR4, GLUT1, GAPDH |
| Fibrosis | FGF2, integrin α5, uPA, PAI1, α1-antitrypsin | FGF7, integrin β1, tenascin C |
| Oncogenesis | HER2, PIM1, EWSR1, CRIP1, NF1, PDCD4, BMI1, MTA2 | KLF4, MALT1, CRK, DMBT1, Daxx, BACH1, MTDH, Ets1, ATM, ZEB1, 14-3-3, BRCA2 |
| Senescence | sirtuin1, sirtuin6, sirtuin7, caveolin1 | klotho, p21 |
| Telomere signaling | TRF1 | TERT |

The nonspecific sequence\*-CEDS, 12A\*-CEDS induced the suppression (blue) and activation (red) of signaling pathways by downregulating (blue) and upregulating (red) the proteins, respectively.

The results indicate that CEDS targeting is variable depending on the context of target sequences and cell types, and simultaneously affects the expression of non-target proteins in the associated pathway and relevant miRNAs. Therefore, it is suggested that the effect of CEDS may be broader and more complicated than we expected.

The present study demonstrated that miR-26a-5p*-CEDS and miR-126-5p*-CEDS induced potent anticancer effect on RAW 264.7 cells by suppressing proliferation-growth-cytodifferentiation signaling, and more the former increased miRNA biogenesis and ER stress signaling, and the latter increased NFkB-immunity signaling, p-53-and FAS-mediated apoptosis signaling. Therefore, it is proposed that the sequential application of miR-26a-5p*-CEDS and miR-126-5p*-CEDS may produce synergistic anticancer effect on RAW 264.7 cells.

Regarding the effect of CEDS on miRNA, CEDS may not only facilitate the hybridization between mature miRNA and target mRNA but also stimulate the miRNA biogenesis by increasing the expression of both primary and mature miRNAs. To know the mechanism of CEDS-induced miRNA biogenesis, further precise molecular genetic study should be required. However, the present study found miR-26a-5p*-CEDS and miR-126-5p*CEDS induced potent anticancer effect on RAW 264.7 cells. Furthermore, since the DNA motifs are quite different from miRNAs in their distributions and functions, the effect of CEDS on DNA motif sequence in genomic DNA will be investigated in the following study.

## Acknowledgments

We would like to express our gratitude to the late Professor Je Geun Chi and the late Dr. Soo Il Chung, who contributed to this research in part.

