## Supplement Fig S1, Table S1-S3 for "Cyclic Electromagnetic DNA Simulation (CEDS) can enhance the Functions of Anticancer miR-26a-5p and miR-126-5p in RAW 264.7 cells"

**The PDF file includes:**

Supplementary Text
Fig. S1.
Tables S1 to S3
References

**Supplementary Text**

***Nonspecific poly-A 12A*-CEDS induced imbalanced protein signaling in RAW 264.7 cells***

The poly-A sequences (4A-12A) are ubiquitously distributed throughout the mouse genome. Therefore, the poly-A sequence may play a role in gene transcription in different regions of the genome, but its function will be variable and nonspecific depending on the context of the genomic environments. In the present study, the poly-A sequence 12A was selected for CEDS experiment as a nonspecific positive control to compare with other CEDS using different miRNA sequences that have been shown to be relevant to gene expression.

RAW 264.7 cells treated with 12A*-CEDS exhibited changes in protein expression throughout essential signaling pathways, both positively and negatively. However, each protein signaling exhibited unique trends of activation or inactivation. As a result, 12A*-CEDS increased protein signaling pathways related to ***proliferation, cMyc/MAX/MAD network, p53/Rb/E2F1 signaling, SHH/PTCH/GLI signaling, Notch/Jagged signaling, miRNA biogenesis, growth factors, cytodifferentiation, RAS signaling, ER stress, angiogenesis, survival, acute inflammation, PARP-mediated apoptosis,*** and ***fibrosis***. On the other hand, it simultaneously activated protein signaling pathways related to ***Wnt/β-catenin signaling, epigenetic modification, protein translation, osteogenesis, neuromuscular differentiation, NFkB signaling, protection, innate immunity, cellular immunity, chronic inflammation, p53-mediated apoptosis, FAS-mediated apoptosis, oncogenesis, senescence, glycolysis***, and ***telomere signaling*** (Fig. S1A).

In RAW264.7 cells treated with 12A*-CEDS, ***proliferation signaling*** appears to be inactivated by downregulating PCNA (up to 90.5%), PLK4 (83.5%), CDK4 (84.2%), nucleolin (80.6%), and upregulating p15/16 (112.5%), p21 (115.7%), and p27 (110.1%), despite the upregulation of Ki-67 (107.4%) and cyclinD2 (106.2%). ***cMyc/MAX/MAD network*** appears to be inactivated by downregulating MAX (86.7%), and upregulating p27 (110.1%).

***p53/Rb/E2F1 signaling*** was inactivated by downregulating Rb1 (86.1%), CDK4 (84.2%), E2F1 (85.9%), as well as coincidentally upregulating p53 (105.8%), E2F4 (136.1%). ***SHH/PTCH/GLI signaling*** was inactivated by downregulating GLI1 (76.1%) despite the slight upregulation of SHH (109%). ***Notch/Jagged signaling*** appears to be inactivated by downregulating Jagged2 (86%) despite the upregulation of Notch1 (119.7%).

***Growth factor signaling pathways*** were partially inactivated by downregulating EGF (73.9%), TGFβ1 (79.3%), TGFβ2 (93.8%), TGFβ3 (74.1%), ERβ (81.2%), SMCD7 (69.7%), FGF2 (93.3%), despite the upregulation of GHRH (116.7%), IGF1 (106.9%),, ALK1 (109.3%), SMAD2/3 (117.5%), HGFα (120.4%), FGF7 (115.4%), progesterone (106.7%). ***Cytodifferentiation signaling*** were partially inactivated by downregulating vimentin (94.5%), TGase2 (87.6%), calnexin (78.7%), CRIP1 (81.6%), AP1M1 (89.1%), caveolin1 (94.7%), DLX2 (89.7%), despite the upregulation of β-actin (110.2%), GAPDH (107.7 %), FAK (106.1%), Ep-CAM (136.9%), VE-cadherin (114.2%), SOX9 (110.6%), tenascin C (108.7%).

***RAS signaling*** appears to be inactivated by downregulating HRAS (82.2%), pAKT1/2/3 (94.1%), RAF-B (80.8%), MEKK (80.6%), ERK1 (86.2%), PTEN (77.2%), STAT3 (86%), PGC1α (93%), PKC (86. 7%), JAK2 (93.8%), p-TYK2 (82.2%), SOS1 (83.6%), despite the upregulation of KRAS (105.1%), PI3K (107.7%), Rab1 (121.7%), p-ERK1 (108.8%), p-STAT3 (115.4%), PKA1α (120.3%), AMPKα (111.9%), p-JNK1 (105.1%), p-PKCα (118.8%).

***ER stress signaling*** was inactivated by downregulating HSP27 (77.5%), eIF2α (84.5%), p-GADD153 (82.5%), BIP (94.4%), despite the upregulation of ATF4 (113.2%), LC3β (105%). ***Angiogenesis signaling*** was suppressed by downregulating HIF1α (94.7%), VEGFA (67%), vWF (86.4%), CD31 (90.3%), CMG2 (78.9%), PDGFA (91.8%), FGF2 (03.3%), despite the upregulation of FLT4 108.9%), LYVE1 (110.7%), angiogenin (115%). ***Survival signaling*** was suppressed by downregulating SP1 (75%), sirtuin1 (80.5%), sirtuin6 (89.6%), sirtuin7 (82.9%), survivin (84.1%), despite the upregulation of SP3 (141.3%).

***Acute inflammation signaling*** appears to be inactivated by downregulating TNFα (93.1%), TRAF6 (83.6%), CXCR4 (85.5%), despite the upregulation of IL8 (122.6%), MCP1 (106.2%), MMP1 (129.2%), CRP (111.7%). ***PARP-mediated apoptosis signaling*** was inactivated by downregulating PARP1 (92.9%), despite the upregulation of AIF1 (108.6%). ***Fibrosis signaling*** was inactivated by downregulating FGF2 (93.3%), integrin α5 (88.5%), uPA (73.6%), PAI1 (81.6%), α1-antitrypsin (88.4%), despite the upregulation of FGF7 (115.4%), integrin β1 (117.5%), tenascin C (108.7%).

Conversely, ***Wnt/β-catenin signaling*** appears to be activated by upregulating Wnt1 (112.1%), β-catenin (112%). ***Protein translation signaling*** appears to be activated by upregulating DHPS (133.5%), eIF5A1 (109%), despite the downregulation of eIF5A2 (90.6%), eIF2α (84.5%). ***Osteogenesis signaling*** was activated by upregulating BMP2 (110.9%), BMP3 (128.4%), BMP4 (114.2%), OPG (106%), RANKL (116.5%), osteonectin (118.5%), aggrecan (111.4%), despite the downregulation of RUNX2 (88.7%), osteopontin (72.7%), osteocalcin (83.4%).

***Neuronal differentiation signaling*** was activated by upregulating NSEγ (106.3%), GFAP (118.9%), despite the downregulation of NK1R (75.4%), while ***cytodifferentiation signaling*** appeared to be inactivated by downregulating myosin-1a (73.9%), MYH2 (87.8%), despite the upregulation of αSMA (119%). ***NFkB signaling*** appears to be activated by upregulating p-p38

(116%), mTOR (113.2%), NFATc1 (119%), NFAT5 (128.9%) and coincidentally upregulating IKK (93.7%), despite the downregulation of MDR (87.3%), GADD45 (91.5%), SRC1 (87.5%).

***Protection signaling*** appears to be inactivated by upregulating SVCT2 (126%), SOD1 (123.5%), HO1 (113.7%), FOXP3 (112.8%), GSTO1/2 (108.7%), NRF2 (108.3%), despite the downregulation of FOXP4 (92.4%), HSP70 (88.8%), FOXO1 (88.6%), hepcidin (86.1%), HSP27 (77.5%). ***Innate immunity signaling*** was activated by upregulating CD56 (114.5%), lactoferrin (134.8%), β-defensin1 (128%), TLR2 (115%), TLR3 (120.2%), TLR4 (116.2%), despite the downregulation of LL37 (75.6%), CAMP (94.1%), α1-antitrypsin (88.4%). ***Celluar immunity signaling*** appears to be activated by upregulating CD3 (up to 114.7%), CD8 (110.6%), CD20 (108.4%), CD44 (105.7%), granzyme B (111.7%), PDL1 (114.1%), despite the downregulation of CD40 (89.8%), CD40 (89.8%), IL28 (68.8%), versican (70.5%), perforin (68%), PD1 (90%).

***Chronic inflammation signaling*** appear to be activated by upregulating IL12 (112%), lysozyme (137.5%), MMP2 (121.5%), MMP9 (119.3%), MMP10 (108.2%), COX1 (122.9%), COX2 (115.9%), LTA4H (125%), despite the downregulation of IL10 (75.3%), M-CSF (91.5%), CD31 (90.9%), CD68 (88.6%), MMP12 (93.9%), TIMP1 (85.5%), TIMP2 (86.3%). ***p53-mediated apoptosis signaling*** was activated by upregulating p53 (105.8%), BAK (118.9%), BAX (108.9%), BCL2 (146%), MDM2 (114.3%), caspase3 (109.1%), c-caspase3 (110.4%), despite the downregulation of p73 (81.6%), APAF1 (84.1%), caspase9 (83.8%). ***FAS-mediated apoptosis signaling*** appears to be activated by upregulating FASL (124.1%), caspase8 (109.4%), caspase3 (109.1%), c-caspase3 (110.4%), despite the downregulation of FAS (80.6%), BID (81.2%).

***Senescence signaling*** appears to be activated by upregulating klotho (142.3%), p21 (115.7%), despite the downregulation of sirtuin1 (82.9%), sirtuin6 (90.3%), sirtuin7 (82.9%), caveolin1 (94.7%). ***Glycolysis signaling*** appears to be activated by upregulating LGR4 (113%), GLUT1 (112.7%), GAPDH (107.7%), despite the downregulation of TIGAR (90.1%), PPARγ (83.9%). ***Telomere signaling*** was activated by upregulating TERT (up to 113.2%), downregulating TRF1 (92.9%).

In addition, 12A*-CEDS markedly influenced ***epigenetic modification signaling*** by downregulating HDAC10 (91.5%), DMAP1 (89.9%), DNMT1 (94.1%), as well as coincidentally upregulating KDM4D (112.2%), MBD4 (106.3%), PCAF (108.1%), HMGB1 (127.8%), BRG1 (115.9%), Ac-lysine (108.5%), lamin A/C (105.5%). Consequently, 12A*-CEDS induced a trend towards an increase in the methylation of histones and DNAs, transcriptional repression.

12A*-CEDS significantly suppressed ***oncogenesis signaling*** by downregulating HER2 (94.5%), PIM1 (89.9%), EWSR1 (86%), CRIP1 (81.6%), NF1 (73.9%), PDCD4 (73.6%), BMI1 (70.6%), MTA2 (67%), while it widely activating ***oncogenesis signaling*** by upregulating KLF4 (135.8%), MALT1 (135.7%), CRK (129.8%), DMBT1 (128.1%), Daxx (126.6%), BACH1 (118.5%), MTDH (116.4%), Ets1 (116.3%), ATM (110.8%), ZEB1 (110.5%), 14-3-3 (109.2%), BRCA2 (107.4%). Among the 28 oncoproteins, 12 were found to be overexpressed, eight were overexpressed, and eight exhibited minimal change in expression compared to the untreated controls.

Consequently, the nonspecific sequence 12A*-CEDS significantly impacted the protein signaling pathways in RAW 264.7 cells. 12*-CEDS suppressed ***RAS signaling***, subsequently inactivating ***proliferation-related signaling axis, ER stress-angiogenesis-survival-acute inflammation signaling axis,*** and ***PARP-mediated apoptosis-fibrosis signaling axis***. In contrast, it enhanced ***NFkB signaling***, subsequently activating ***p53- and FAS-mediated apoptosis***

***signaling axis,*** and ***senescence-glycolysis-oncogenesis-innate immunity-chronic inflammation signaling axis*** (Fig. S1B, Table S3).

The results indicate that 12*-CEDS also affects the protein signaling pathways in a similar manner to the above experiments, due to the existence of numerous poly-A sequences (4A-12A) in the mouse genome. 12A*-CEDS resulted in imbalanced protein signaling pathways in RAW 264.7 cells, activating epigenetic methylation, oncogenesis, and telomere instability, leading to chronic inflammation and apoptosis, in the absence of cellular growth and differentiation, wound healing, and ROS protection. Therefore, it is postulated that 12A*-CEDS may influence the cells to undergo aging and retrogressive changes with a high potential of oncogenesis.

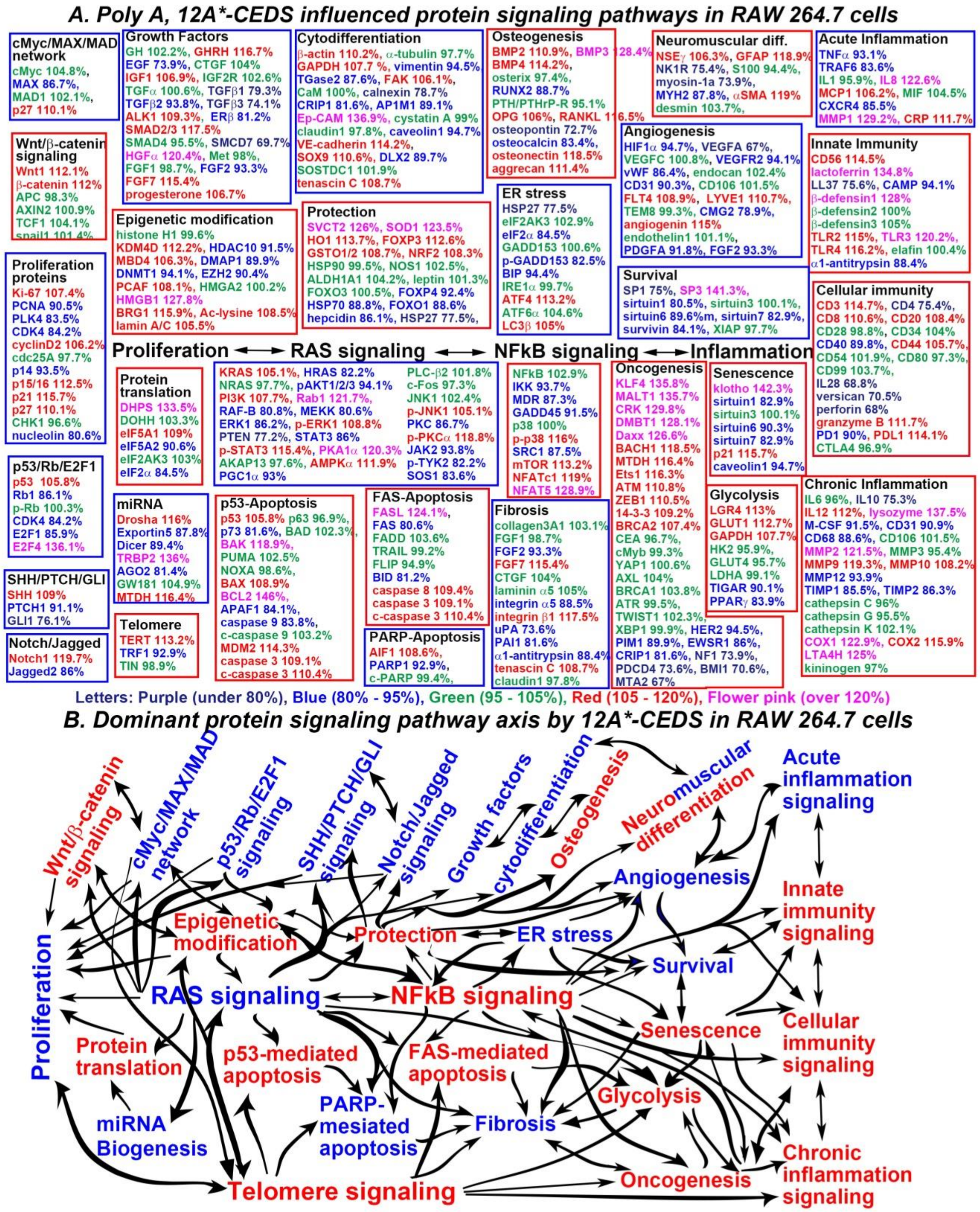


**Figure S1.** The nonspecific sequence 12A*-CEDS influenced the protein signaling pathways (A) and axis (B) in RAW 264.7 cells. The IP-HPLC results revealed proteins that were downregulated (blue), upregulated (red), and minimally changed (green) compared to the untreated controls. This allowed the dominant trend of suppressed (blue square) and activated (red square) signaling pathways to be defined.

**Table S1.** The primer pairs of miRNAs used in quantitative real time PCR (qPCR)

| mature mmu-miRNA | primer direction | mature miRNA primer sequences | Primary miRNA primer sequences |
|---|---|---|---|
| **miR-26a-5p** | Forward | TTCAAGTAACC | AAGCTGGAGGACCGA |
| | Reverse | AGCCTATCC | ACTCTGTTGTTGCCG |
| **miR-34a-5p** | Forward | TAGGCAGTG | CCCGGTGCTCGGTTT |
| | Reverse | GTACAATCA | GCGGGTGATGCTGTG |
| **miR-126-5p** | Forward | CATTATTAC | ACAGCGCGTACC |
| | Reverse | GCGCGTACC | GAGTGCCAGCCGTGG |
| **miR-150-5p** | Forward | CCTGTCTCCCA | CAGCACTGGTAC |
| | Reverse | CACTGGTAC | CTGGAGTTGTTGCTG |
| **miR-155-5p** | Forward | TTAATGCTA | TTAATGCTA |
| | Reverse | AACCCCTA | CCATTTGTTCCATGT |
| **miR-181a-5p** | Forward | TACAATCAACG | TACAATCAACG |
| | Reverse | ACTCACCGAC | TAGGGTACAATCAAC |
| **miR-216a-5p** | Forward | TAATCTCAG | TAATCTCAG |
| | Reverse | TCACAGTTGCC | TTAGCATAATCCCAG |
| **miR-365a-3p** | Forward | TAAGGATTT | ATCTGCCCCT |
| | Reverse | ATCTGCCCCT | TATCTTACAGAGTTG |
| **miR-655-3p** | Forward | AAAGAGGTTA | AAAGAGGTTA |
| | Reverse | GAACATAACACG | AACTATGCAAGG |

**Table S2.** Antibodies used in the IP-HPLC analysis.

| Signaling | No. | Antibodies |
|---|---|---|
| **Proliferation** | 12 | Ki-67, PCNA, PLK4, CDK4, cyclin D2, cdc25A, p14, p15/16, p21, p27, CHK1, nucleolin |
| **cMyc/MAX/MAD network** | 4 (1) | cMyc, MAX, MAD1, (p27) |
| **Wnt/β-catenin signaling** | 6 | Wnt1, β-catenin, APC, AXIN2, TCF1, Snail |
| **p53/Rb signaling** | 6(1) | p53, Rb1[#], p-Rb, (CDK4), E2F1, E2F4 |
| **SHH/PTCH/GL1 and Notch/Jagged signaling** | 5 | SHH, PTCH, GL1, Notch1, Jagged2 |
| **Epigenetic modification** | 13 | Histone H1, KDM4D[$], HDAC10[$], MBD4, DMAP1, DNMT1, EZH2, PCAF, HMGA2, HMGB1, BRG1, Ac-lysine, lamin A/C |
| **Protein translation** | 6 | DOHH[!], DHPS[!], eIF5A1[!], eIF5A2[!], eIF2AK3, eIF2α^ |
| **miRNA biogenesis** | 7 | Drosha, Exportin5, Dicer, TRBP2, AGO2, GW182, MTDH |
| **Growth factors** | 22 | GH, GHRH, EGF, EGFR, IGF1, IGF2R, TGFα, TGFβ1[$], TGFβ2, TGFβ3, ALK1, SMAD2/3, SMAD4, SMAD7, HGFα, Met, FGF1, FGF2, FGF7, CTGF, ERβ, progesterone |
| **RAS signaling** | 26 | KRAS[$], HRAS, NRAS[$], pAKT1, PI3K, Rab1, RAF-B, MEKK, ERK1, p-ERK1[$], PTEN[&], STAT3, p-STAT3, PKA1α, AKAP13, AMPKα[@], PGC1α, PLCβ2, c-Fos, JNK1, p-JNK1, PKC, p-PKC1α[@], p-TYK2, JAK2[$], SOS1 |
| **NFkB signaling** | 10 | NFkB, IKK, MDR, GADD45, p38, p-p38, SRC1, mTOR[@], NFATc1, NFAT5 |
| **p53-mediated apoptosis** | 18 (1) | (p53), p63, p73, MDM2, BAD, BAK, PUMA, NOXA, BAX, APAF1, BCL2, caspase9, c-caspase9, AIF1, PARP1, c-PARP, caspase3, c-caspase3 |
| **FAS-mediated apoptosis** | 9 | FASL, FAS, FADD, TRAIL, FLIP, BID, caspase8, caspase3, c-caspase3 |
| **Cytodifferentiation** | 19 | β-actin, α-tubulin, GAPDH, vimentin, TGase2[+], FAK, CaM, calnexin, CRIP1, AP1M1, Ep-CAM, cystatin A, claudin1, caveolin1, VE-cadherin[&], SOX9, DLX2, SOSTDC1, tenascin C |
| **Osteogenesis** | 11 | BMP2, BMP3, BMP4, RUNX2, osterix, OPG, RANKL, osteopontin, osteonectin, osteocalcin, aggrecan |
| **Neuromuscular differentiation** | 8 | NSEγ, GFAP, NK1R, S100, myosin1a, MYH2, desmin, αSMA |
| **Protection** | 15 | HSP27, HSP70, HSP90, SOD1, GSTO1/2, NOS1, HO1, NRF2, hepcidin, SVCT2, leptin, FOXO1, FOXO3, FOXP3, FOXP4 |
| **ER stress** | 10 | HSP27, eIF2AK3, eIF2α^, GADD153, p-GADD153, BIP, IRE1α, ATF4, ATF6α, LC3β, |
| **Angiogenesis** | 16 | HIF1α[&], VEGFA, VEGFC, VEGFR2, vWF[$], endocan, CD31, CD106, FLT4[$], LYVE1, TEM8, CMG2[$], angiogenin, endothelin1, PDGFA, FGF2 |
| **Survival** | 8 | SP1[@], SP3[@], sirtuin1, sirtuin3, sirtuin6, sirtuin7, survivin[@], XIAP |
| **Fibrosis** | 11 (2) | collagen3A1, FGF1, CTGF, laminin α5, integrin α5, integrin β1, uPA[+], PAI1, α1-antitrypsin, (tenascin C, claudin1) |
| **Oncogenesis** | 29 (1) | CEA[$], cMyb, 14-3-3, YAP1, DMBT1, MTDH, CRK, PIM1, AXL, BACH1, BMI1, KLF4, ZEB1, HER2, BRCA1[&], BRCA2[&], NF1, ATR, ATM, PDCD4, CRIP1, TWIST1, XBP1, MTA2, EWSR1, Ets1, Daxx, MALT1, (CRIP1) |
| **Senescence** | 7 (2) | Klotho, sirtuin1, sirtuin3, sirtuin6, sirtuin7, (p21, caveolin1) |
| **Telomere** | 3 | TERT, TRF1, TIN2 |
| **Metabolism** | 8 | HX2, GAPDH, GLUT1, GLUt4, LGR4, PPARγ, TIGAR, LDHA |
| **Acute inflammation** | 9 | TNFα[@], TRAF6, IL1, IL8, MCP1, MIF, CXCR4, MMP1, CRP |
| **Innate immunity** | 12 | CD56, elafin, lactoferrin, LL37, CAMP, β-defensin 1, -2, -3, TLR2, TRL3, TLR4, α1-antitrypsin[&] |
| **Cellular immunity** | 19 | CD3, CD4, CD8, CD20, CD28, CD34, CD40, CD44, CD54, CD80, CD99, IL28, versican, perforin, granzyme B, PD1, PDL1, CTLA4, LTA4H |
| **Chronic inflammation** | 21 (2) | IL6, IL10, IL12, lysozyme, M-CSF, (CD31), CD68, (CD106), MMP2, MMP3, MMP9, MMP10, MMP12, TIMP1, TIMP2, cathepsin C, cathepsin G, cathepsin K, COX1, COX2, kininogen |
| **Total** | **350 (10)** | |

( ): Overlapped antibodies.

[#] DAKO, Denmark; [$] Neomarkers, CA, USA; [@] ZYMED, CA, USA; [&]Abcam, Cambridge, UK; ^Cell signaling technology, USA; [!] kindly donated from M. H. Park in NIH, USA (*1*); [+] kindly donated from S. I. Chung in NIH, USA (*2, 3*); no mark; Santa Cruz Biotechnology, USA; the number of antibodies overlapped; ( ).

**Abbreviations:** 14-3-3: the 14th fraction eluting from a DEAE-cellulose column and in position 3.3 on a starch electrophoresis gel, Ac-lysine: Acetylated-Lysine, AGO2: argonaute 1, Eukaryotic translation initiation factor 2C (eIF2C), AIF1: apoptosis inducing factor 1, AKAP13: A-kinase anchoring proteins 13, ALK1: activin receptor-like kinase 1, AMPKα: AMP-activated protein kinase α, pAKT: v-akt murine thymoma viral oncogene homolog, p-Akt1/2/3 phosphorylated (p-Akt, Thr 308), AP1M1: AP-1 complex subunit mu-1, APAF1: apoptotic protease-activating factor 1, APC: adenomatous polyposis coli, ATF4: activating transcription factor 4, ATM: ataxia telangiectasia caused by mutations, ATR: ataxia telangiectasia and Rad3-related protein, AXIN2: axis inhibition protein 2, AXL: Tyrosine-protein kinase receptor UFO, BACH1: Transcription regulator protein, BAD: BCL2 associated death promoter, BAK: BCL2 antagonist/killer, BAX: BCL2 associated X, BCL2: B-cell leukemia/lymphoma-2, BID: BH3 interacting-domain death agonist, BIP (GRP 78): binding immunoglobulin protein, BMI1: B lymphoma Mo-MLV insertion region 1 homolog, polycomb group RING finger protein 4, BMP2: bone morphogenesis protein 2, BRCA1: breast cancer type 1 susceptibility protein, BRG1 (SMARCA4): transcription activator, c-caspase 3: cleaved-caspase 3, CaM: calmodulin, CAMP: Cathelicidin antimicrobial peptide, CD3: cluster of differentiation 3, CD31: Platelet endothelial cell adhesion molecule (PECAM-1), CD44 (HCAM): homing cell adhesion molecule, CD54 (ICAM1): intercellular adhesion molecule 1, CD56 (NCAM): neural cell adhesion molecule 1, CD106 (VCAM1): vascular cell adhesion molecule-1, cdc25A: cell division cycle 25A, CDK4: cyclin dependent kinase 4, CEA: carcinoembryonic antigen, CHK1: Checkpoint kinase 1, CMG2: capillary morphogenesis protein 2 (anthrax toxin receptor 2), COX1: cyclooxygenase-2, CRIP1: Cysteine-rich intestinal protein 1, CRK: proto-oncogene c-Crk, CRP: C-reactive protein, CTGF: connective tissue growth factor, CTLA4: cytotoxic T lymphocyte-associatedprotein-4, CXCR4: C-X-C chemokine receptor type 4, Daxx: Death-associated protein 6, DHS: deoxyhypusine synthase, DLX2, homeobox protein Distal-less (Dlx) family, DMAP1: DNA methyltransferase 1 associated protein, DMBT1: deleted in malignant brain tumors 1, DNMT1: DNA 5-cytosine methyltransferase 1, DOHH: deoxyhypusine hydroxylase, E2F1: transcription factor, eIF2AK3 (PERK): protein kinase R (PKR)-like endoplasmic reticulum kinase (PERK), EGF: Epidermal growth factor, EGF: Epidermal growth factor receptor, elF5A1: eukaryotic translation initiation factor 5A-1, EpCAM: Epithelial cell adhesion molecule, ERβ: estrogen receptor beta, ERK1 (extracellular signal-regulated protein kinase 1, MAPK3 (mitogen-activated protein kinase 3)), Ets1: Protein C-ets-1, EZH2 (ENX-1): enhancer of zeste homolog 2, EWSR1: Ewing Sarcoma Breakpoint Region 1 Protein, FADD: FAS associated via death domain, FAK: focal adhesion kinase, FAS: CD95/Apo1, FASL: FAS ligand, FGF1: fibroblast growth factor 1, FLIP: FLICE-like inhibitory protein, FLT4: Fms-related tyrosine kinase 4, FOXO1: Forkhead/winged helix box gene, group O 1, FOXP3: forkhead box P3, GADD45: growth arrest and DNA-damage-inducible 45, GADD153: C/EBP homologous protein (CHOP), GAPDH: glyceraldehyde 3-phosphate dehydrogenase, GFAP: glial fibrillary acidic protein, GH: growth hormone, GHRH: growth hormone-releasing hormone, GL1: Glioma-associated oncogene 1, GLUT1: Glucose transporter 1, GSTO1/2: glutathione S-transferase ω 1/2, GW182: Trinucleotide repeat-containing gene 6A protein, HDAC10: histone deacetylase 10, HER2: EGF receptor tyrosine-protein kinase erbB-2, HGFα: hepatocyte growth factor α, HIF1α: hypoxia inducible factor-1α, HO-1: heme oxygenase 1, HMGA2: High mobility group (HMG) proteins A2, HRAS: GTPase HRAS, HSP70: heat shock protein 70, HX2: hexokinases 2, IGF1: insulin-like growth factor 1, IGF2R: insulin-like growth factor 2 receptor, IKK: ikappaB kinase, IL1: interleukin-1, IRE1α (ERN1): inositol-requiring enzyme 1 α, JAK2: Janus kinase 2, JNK1: Jun N-terminal protein kinase 1, KDM4D: Lysine-specific demethylase 4D, KLF4: Kruppel-like factor 4, KRAS: V-Ki-ras2 Kirsten rat sarcoma viral oncogene homolog, LC3β: microtubule-associated protein 1A/1B-light chain 3β, LDHA: Lactate dehydrogenase A, LGR4: Leucine-rich repeat-containing G-protein coupled receptor 4, LTA4H: leukotriene A4 hydrolase, LYVE1: lymphatic vessel endothelial hyaluronan receptor 1, MAD1: mitotic arrest deficient 1, MALT1: Mucosa-associated lymphoid tissue lymphoma translocation protein 1, MAX: myc-associated factor X, MBD4: methyl-CpG-binding domain protein 4, MCP1: monocyte chemotactic protein 1, M-CSF: macrophage colony-stimulating factor, MDM2: mouse double minute 2 homolog, MDR: multiple drug resistance, MEKK1, MMP1: matrix metalloprotease 1, Met: tyrosine-protein kinase Met, MIF: Macrophage migration inhibitory factor, MMP1: matrix metalloproteinase 1, MTA2: Metastasis-associated protein 2, MTDH: Metadherin, mTOR: mammalian target of rapamycin, cMyc: V-myc myelocytomatosis viral oncogene homolog, MYH2: myosin-2, NF1: neurofibromin 1, NFAT5: nuclear factor of activated T cells 5, NFATc1: nuclear factor of activated T-cells, cytoplasmic 1, NFkB: nuclear factor kappa-light-chain-enhancer of activated B cells, NK1R: neurokinin 1 receptor, NOS1: nitric oxide synthase 1, Notch1: Notch homolog 1, NOXA: Phorbol-12-myristate-13-acetate-induced protein 1, NRAS: neuroblastoma RAS Viral Oncogene homolog, NRF2: nuclear factor (erythroid-derived)-like 2, NSEγ: Neuron specific γ enolase γ, OPG: osteoprotegerin, PAI-1: plasminogen activator inhibitor-1, PARP-1: poly-ADP ribose polymerase 1, c-PARP-1: cleaved-PARP-1, PCAF: p300/CBP-associated factor, PCNA: proliferating cell nuclear antigen, PD-1 (CD279): programmed cell death protein 1, PDCD4: Programmed cell death protein 4, PDGFA: platelet-derived growth factor-A, PDL1: Programmed death-ligand 1, PGC-1α: peroxisome proliferator-activated receptor gamma coactivator 1α, PI3K: phosphatidylinositol-3-kinase, PIM-1: Proto-oncogene serine/threonine-protein kinase 1, PKA1α: protein kinase A 1α, PKC: protein kinase C, PLCβ2: 1-phosphatidylinositol-4,5-bisphosphate phosphodiesterse β-2, PLK4: polo like kinase 4 or serine/threonine-protein kinase, PPARγ: Peroxisome proliferator- activated receptor gamma, PTCH: Protein patched homolog 1, PTEN: phosphatase and tensin homolog, PUMA: p53 upregulated modulator of apoptosis, Rab1: RAS-related protein, Rab GTPases, RAF-B: v-Raf murine sarcoma viral oncogene homolog B, RANKL: receptor activator of nuclear factor kappa-B ligand, Rb-1: retinoblastoma-1, RUNX2: Runt-related transcription factor-2, SHH: sonic hedgehog, sirtuin3: silent mating type information regulation 2 homolog 3, NAD-dependent deacetylase, α-SMA: alpha-smooth muscle actin, SMAD4: mothers against decapentaplegic, drosophila homolog 4, SOD1: superoxide dismutase-1, SOS1: son of sevenless homolog 1, SOSTDC1: sclerostin domain-containing protein 1**,** SOX9: SRY (sex-determining region Y)-related HMG-box transcription factor 9, SP1: specificity protein 1, SRC1: steroid receptor coactivator-1, STAT3: signal transducer and activator of transcription-3, SVCT2: sodium-dependent vitamin C transporter 2, TCF1: T-cell factor-1, TEM8: tumor-specific endothelial marker 8 (anthrax toxin receptor 1), TERT: human telomerase reverse transcriptase, TGase 2: transglutaminase 2, TGF-β1: transforming growth factor-β1, TIGAR: TP53-induced glycolysis and apoptosis regulator, TIMP1: Tissue inhibitor of metalloproteinases 1, TIN2: TERF1-interacting nuclear factor 2, TLR2: toll-like receptor 2, TNFα: tumor necrosis factor-α, TRAF6: TNF receptor associated factor 6, TRAIL: TNF-related apoptosis-inducing ligand, TRBP2, trans-activation-responsive (HIV-1) RNA binding protein 2, TRF1: Telomeric Repeat Factor 1, TWIST1: Twist-related protein 1, p-TYK2: Non-receptor tyrosine-protein kinase, uPA: urokinase-type plasminogen activator, VE-cadherin: vascular endothelial cadherin, VEGFA vascular endothelial growth factor A, VEGFR2: vascular endothelial growth factor receptor 2, p-VEGFR2: phosphorylated vascular endothelial growth factor receptor 2 (Y951), vWF: von Willebrand factor, Wnt1: proto-oncogene protein Wnt1, XBP1: X-box binding protein 1, XIAP: X-linked inhibitor of apoptosis protein, YAP1: Yes-associated protein 1, ZEB1: Zinc finger E-box-binding homeobox 1

**Table S3.** Table 17. Protein expressions in RAW 264.7 cells treated with nonspecific sequence 12A*-CEDS

| Signaling pathways | Downregulated Proteins | Upregulated Proteins |
|---|---|---|
| Proliferation | PCNA, PLK4, CDK4, p14, nucleolin | Ki-67, cyclin D2, p15/16, p21, p27 |
| cMyc/MAX/MAD network | MAX | p27 |
| Wnt/β-catenin signaling | | Wnt1, β-catenin |
| p53/Rb/E2F1 signaling | Rb1, CDK4, E2F1 | p53, E2F4 |
| SHH/PTCH/GLI signaling | PTCH1, GLI1 | SHH |
| Notch/Jagged signaling | Jagged2 | Notch1 |
| Epigenetic modification | HDAC10, DMAP1, DNMT1, EZH2 | KDM4D, MBD4, PCAF, HMGB1, BRG1, Ac-lysine, lamin A/C |
| Protein translation | eIF5A2, eIF2α | DHPS, eIF5A1 |
| miRNA biogenesis | Exportin5, Dicer, AGO2, | Drosha, TRBP, MTDH |
| Growth factors | EGF, TGFβ1, TGFβ2, TGFβ3, SMCD7, FGF2, ERβ | GHRH, NF1, ALK1, SMCD2/3, HGFα, FGF7, progesterone |
| Cytodifferentiation | Vimentin, TGase2, calnexin, CRIP1, AP1M1, caveolion1, DLX2 | β-actin, GAPDH, FAK, Ep-CAM, VE-cadherin, SOX9, tenascin C |
| Osteogenesis | RUNX2, osteopontin, osteocalcin, | BMP2, BMP3, BMP4, OPG, RANKL, osteonectin, aggrecan |
| Neuromuscular differentiation | NK1R, myosin-1a, MYH2 | NSEγ, GFAP, αSMA |
| RAS signaling | HRAS, pAKT1/2/3, RAF-B, MEKK, ERK1, PTEN, STAT3, PGC1α, PKC, JAK2, p-TYK2, SOS1 | KRAS, PI3K, Rab1, p-STAT3, PKA1α, AMPKα, p-JNK1, p-PKC1α |
| NFkB signaling | IKK, MDR, GADD45, SRC1 | p-p38, mTOR, NFATc1, NFAT5 |
| p53-mediated apoptosis | p73, APAF1, caspase9 | p53, BAK, BAX, BCL2, MDM2, caspase3, c-caspase3 |
| FAS-mediated apoptosis | FAS, BID | FASL, caspase8, caspase3, c-caspase3 |
| PARP-mediated apoptosis | PARP1 | AIF1 |
| Protection | FOXP4, HSP70, FOXO1, hepcidin, HSP27 | SVCT2, SOD1, HO1, FOXP3, GSTO1/2, NRF2, |
| ER stress | HSP27, eIF2α, p-GADD153, BIP | ATF4, LC3β |
| Angiogenesis | HIF1α, VEGFA, VEGFR2, vWF, CD31, CMG2, PDGFA, FGF2 | FLT4, LYVE1, angiogenin |
| Survival | SP1, sirtuin1, sirtuin1, sirtuin6, sirtuin7, survivin | SP3 |
| Acute inflammation | TNFα, TRAF6, CXCR4 | IL8, MCP1, MMP1, CRP |
| Innate immunity | LL37, CAMP, α1-antitrypsin | CD56, lactoferrin, TLR2, TLR3, TLR4 |
| Cellular immunity | CD4, CD40, IL28, versican, perforin, PD1 | CD3, CD8, CD20, CD44, granzyme B, PDL1 |
| Chronic inflammation | IL6, IL10, M-CSF, CD31, CD68, MMP12, TIMP1, TIMP2 | IL12, lysozyme, MMP2, MMP9, MMP10, COX1, COX2, LTA4H |
| Glycolysis | TIGAR, PPARγ | LGR4, GLUT1, GAPDH |
| Fibrosis | FGF2, integrin α5, uPA, PAI1, α1-antitrypsin | FGF7, integrin β1, tenascin C |
| Oncogenesis | HER2, PIM1, EWSR1, CRIP1, NF1, PDCD4, BMI1, MTA2 | KLF4, MALT1, CRK, DMBT1, Daxx, BACH1, MTDH, Ets1, ATM, ZEB1, 14-3-3, BRCA2 |
| Senescence | sirtuin1, sirtuin6, sirtuin7, caveolin1 | klotho, p21 |
| Telomere signaling | TRF1 | TERT |

The nonspecific sequence*-CEDS, 12A*-CEDS induced the suppression (**blue**) and activation (**red**) of signaling pathways by downregulating (**blue**) and upregulating (**red**) the proteins, respectively.
